# Single-cell reference mapping to construct and extend cell-type hierarchies

**DOI:** 10.1101/2022.07.07.499109

**Authors:** Lieke Michielsen, Mohammad Lotfollahi, Daniel Strobl, Lisa Sikkema, Marcel J.T. Reinders, Fabian J. Theis, Ahmed Mahfouz

**Affiliations:** Department of Human Genetics, Leiden University Medical Center, Einthovenweg 20, 2333ZC, Leiden, The Netherlands; Leiden Computational Biology Center, Leiden University Medical Center, Einthovenweg 20, 2333ZC, Leiden, The Netherlands; Delft Bioinformatics Lab, Delft University of Technology, Van Mourik Broekmanweg 6, 2628XE, Delft, The Netherlands; Institute of Computational Biology, Helmholtz Zentrum München, Munich, Germany; Institute of Clinical Chemistry and Pathobiochemistry, TUM School of Medicine, Technical University of Munich, 81675 Munich, Germany; TUM School of Life Sciences Weihenstephan, Technical University of Munich, Germany; Department of Mathematics, Technical University of Munich, Munich, Germany

**Author notes:** Correspondence to &. These authors contributed equally (Lieke Michielsen, Mohammad Lotfollahi).

## Abstract

Single-cell genomics is now producing an ever-increasing amount of datasets that, when integrated, could provide large-scale reference atlases of tissue in health and disease. Such atlases increase the scale and generalizability of analyses and enable combining knowledge generated by individual studies. Specifically, individual studies often differ regarding cell annotation terminology and depth, with different groups often using distinct terminology. Understanding how annotations are related and complement each other would mark a major step towards a consensus-based cell-type annotation reflecting the latest knowledge. Whereas recent computational techniques, referred to as “reference mapping” methods, facilitate the usage and expansion of existing reference atlases by mapping new datasets (i.e., queries) onto an atlas; a systematic approach towards harmonizing dataset-specific cell-type terminology and annotation depth is still lacking. Here, we present “treeArches”, a framework to automatically build and extend reference atlases while enriching them with an updatable hierarchy of cell-type annotations across different datasets. We demonstrate various use cases, from automatically resolving relations between reference and query cell types to identifying unseen cell types absent in the reference, such as disease-associated cell states. We envision treeArches enabling data-driven construction of consensus atlas-level cell-type hierarchies and facilitating efficient usage of reference atlases.

## Introduction

Single-cell sequencing technologies have revolutionized our understanding of human health. Hereto, large single-cell datasets - referred to as “reference atlases” - have been built to characterize the cellular heterogeneity of whole organs. An example is all the organ- and body-scale cell atlases constructed within big consortia such as the human cell atlas (HCA) (1–5). Users can contextualize their datasets within these references to identify novel cell types. This enables the discovery of disease-affected cell types that can be prioritized for treatment design (6–8).

To create a reference atlas, one would ideally leverage information from multiple scRNA-seq datasets and harmonize their cell annotations. This, however, is not as easy as it seems since all datasets are annotated at a different resolution. Furthermore, matching cell types based on their names is difficult. Databases such as ‘Cell Ontology’ try to overcome this problem, but a complete naming convention is still missing (9). When constructing the Human Lung Cell Atlas (HLCA), for instance, the cell type labels of 14 datasets had to be manually harmonized, which is a time-consuming process (2). To accelerate the construction of reference atlases, we developed scHPL: a method to automatically match the cell-type labels of multiple datasets and construct a cell-type hierarchy (10). In follow-up, Novella-Rausell et al. showed how scHPL simplified the process when building a mouse kidney atlas (11).

The concept of a “reference atlas”, however, suggests it should help analyze and interpret new datasets (here denoted as “query”). This is, however, complicated by batch effects between the reference and query, limited computational resources, and data privacy and sharing. Recently, we, along with others, developed computational approaches (known as “reference mapping” methods) to address these challenges (4, 12, 13). Such methods could for instance be used to map a query dataset to the reference and annotate the cells. Currently, there is no method available that tackles both challenges simultaneously.

To address these challenges, we present treeArches, a framework that builds upon single-cell architectural surgery (scArches) (12) and single-cell Hierarchical Progressive Learning (scHPL) (10) to progressively build and update a reference atlas and corresponding hierarchical classifier. Our approach allows users to build a reference atlas using existing integration methods supported by scArches (e.g., scVI, scANVI, totalVI, and all others described in (14)). Next, we use scHPL to augment this reference atlas by learning the relations between cell types to construct a cell-type hierarchy. Afterward, query data, which can be either annotated or unannotated, can be mapped to the reference. If the query is annotated, the query cells can expand the newly updated tree by highlighting potential novel cell types and their relationship with other cell types in the reference. Otherwise, the created reference can be used to annotate the query cells and identify new unseen cell types in the query. Unlike existing methods, we show that treeArches can be used to create a reference atlas and corresponding cell-type hierarchy from scratch, update an existing reference atlas and the hierarchy by finding novel relations between cell types, and leverage a reference atlas to transfer labels to a new dataset.

## Methods

### Overview

treeArches consists of two main steps: (i) removing the batch effects between datasets and (ii) matching the annotated cell types to construct a cell-type hierarchy (Fig 1). Starting with multiple labeled datasets, hereafter called reference datasets, we first use neural network-based reference-building models (e.g., sc(AN)VI (14) or scGen (15)), which are top performers in recent data benchmarking efforts (16) and compatible with scArches, to construct a latent space. Next, we use scHPL to construct the cell-type hierarchy (Fig 1A). For each dataset, we train a classifier in the learned latent space and cross-predict the labels of the other dataset(s). Using the confusion matrices, we automatically match the cell types to create a hierarchy. This hierarchy also represents a hierarchical classifier where every node represents a cell type in one or more of the datasets. Afterwards, we can map new query datasets to the learned latent space using architectural surgery, a transfer learning approach to map query datasets to references, implemented by scArches (Fig 1B). Architectural surgery brings the advantage that the count matrices of the reference datasets are not needed anymore for querying the model. Instead, we only use the pre-trained neural network architecture. The query datasets can either be labeled or unlabeled. In the case of a labeled dataset, we match the cell types from the query to the reference and again update the hierarchy we had learned on the reference datasets. In the case of an unlabeled query, we annotate the cells using the learned hierarchy.

**Figure 1.**
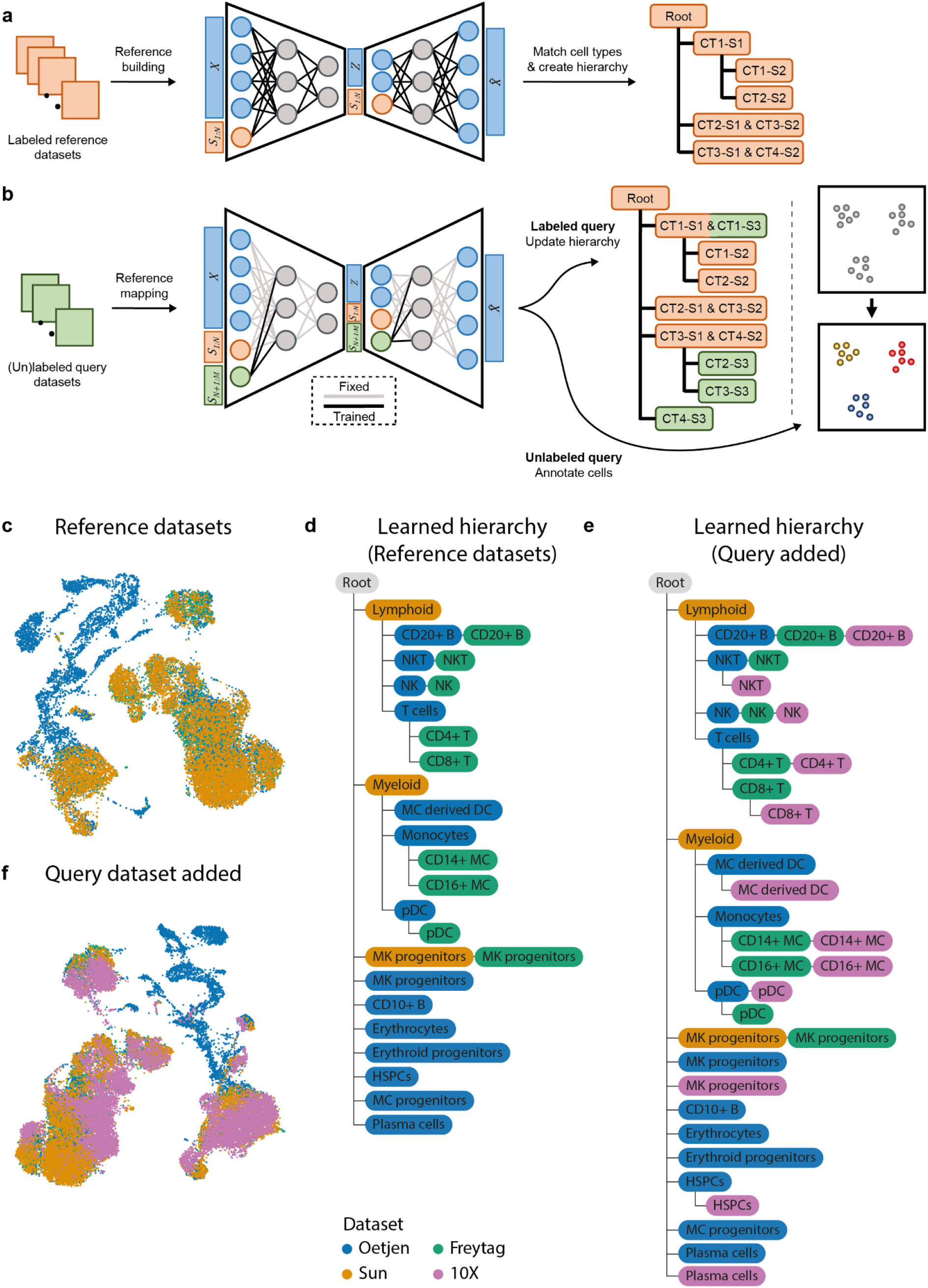
A schematic version of treeArches and an example using PBMC and bone marrow datasets. **(a)** Pre-training of a latent representation using labeled public reference datasets. After integration, a cell-type hierarchy is created by matching the cell types of the different datasets. Here, for instance, cell types (CT) 1 and 2 from study (S) 2 are subtypes of CT1 from S1. **(b)** (Un)labeled query datasets can be added to the latent representation by applying architectural surgery. After integration, the cell-type hierarchy is updated with labeled query datasets. Unlabeled query datasets can be annotated using the learned hierarchy. **(c)** UMAP embedding showing the integrated latent space of the three reference datasets. **(d)** Cell-type hierarchy learned from the three reference datasets. MC derived DC: monocyte-derived dendritic cells, MC: monocytes, pDC: plasmacytoid dendritic cells, HSPC: hematopoietic stem and progenitor cell **(e)** Updated hierarchy after the 10X dataset was added. **(f)** UMAP embedding showing the integrated latent space of the reference and query datasets.

When matching the cell types or predicting labels of a query dataset, it is important to identify new cell types that are not present in the reference. This is only possible when biological variation is preserved when mapping the datasets to the latent space and when the classifier in scHPL recognizes unseen cells, i.e. cells that are not present in the tree. Therefore scHPL adopts a rejection strategy, which rejects these unseen cells and identifies them as a new cell type. Within scHPL, a cell is rejected if it meets one of the following criteria: 1) if the posterior probability of the classifier is lower than a threshold which means the predicted label is ambiguous, 2) if the distance between a cell and its closest neighbors is too big, and 3) if the reconstruction error (when mapping to a reduced PCA space and back) is above a threshold, which means the query cell is too different from the reference cell types. These three thresholds are automatically set based on the distribution of the data.

treeArches is a framework built around scArches (version 0.5.3) (12) and scHPL (version 1.0.1) (10). A detailed description of scArches and scHPL can be found in their original papers (10, 12). Here, we only describe changes to the original methods when combined in the treeArches framework. We enhanced the original version of scHPL by adding the option to use a *k-*nearest neighbor (kNN) classifier. The dimensionality of the latent space learned by scArches is relatively low (varying between 10 and 30 dimensions). We noticed that the linear SVM originally implemented doesn’t perform well, since the cell types are not linearly separable anymore. Therefore, it is better to use scHPL with the kNN classifier in this case. In contrast to the linear SVM, we train a multiclass classifier for every parent node instead of a binary classifier for every child node (10). During training, we set the default number of neighbors to 50. However, when there are cell types in the dataset that consist of less than 50 cells, this is not ideal. Therefore, we added an extra option (*dynamic_neighbors*) to automatically decrease *k* to the size of the smallest cell type across the direct child nodes. Since the tree consists of multiple classifiers, it can thus be that they all use a different number of neighbors because of this option. For the kNN classifier itself, we implemented alternatives using either the FAISS library (17) or the scikit-learn library (18). The FAISS implementation is faster than the scikit-learn library but is only available on Linux.

### Detecting new or diseased cell types

We have implemented three methods to detect new or diseased cell types: 1) a threshold on the posterior probability, 2) a threshold on the reconstruction error, and 3) a threshold on the distance between query and reference. The first two options were already implemented in the previous version of scHPL. The default threshold for the first option is 0.5. The threshold for the second rejection option is determined using a nested cross-validation loop. It is the median reconstruction error that gives a certain amount of false negatives on the test folds (default = 0.5%). The third option rejects cells whose distance to the predicted class is too big. The threshold for rejection is determined by calculating the neighbors for all cells in the training set, averaging the distance across the neighbors, and taking the 99th percentile.

## Datasets

### PBMC datasets

The dataset was obtained from the recent data integration benchmark (16). The data contains bone marrow samples from Oetjen et al. (19) and also PBMC samples that were obtained from 10x Genomics https://support.10xgenomics.com/single-cell-gene-expression/datasets/3.0.0/pbmc_10k_v3, Freytag et al. and Sun et al. (20, 21), the original url and the preprocessing and annotation details can be found in Luecken et al. (16). Marker genes specific to early erythrocytes and platelets were downloaded from Azimuth (4).

### Brain datasets

We used datasets from the primary motor cortex of three species: human, mouse, and marmoset (22). We downloaded the datasets from the Cytosplore comparison viewer. In these datasets, genes were already matched based on one-to-one homologs. For the analysis, we only kept these one-to-one matches (15,860 genes in total). We selected 2,000 highly variable genes based on the reference datasets (mouse and marmoset) and used those counts as input for treeArches. The datasets are annotated at three different resolutions: Class, Subclass, and RNA_cluster. The class level contains three broad brain cell types: GABAergic neurons, glutamatergic neurons, and non-neuronal cells. At the subclass level, the cells are annotated at a higher resolution (5-10 subclasses per class). The RNA_cluster level contains the highest resolution. Here, we will use the subclass level to match the cell types. Marker genes used for visualization were chosen based on Supplementary Tables 5 and 6 from the original paper (22).

### Human Lung Cell Atlas

The human lung cell atlas (HLCA) is a carefully constructed reference atlas for the human respiratory system (2). Sikkema et al. aligned 14 datasets, harmonized the annotations, and built a cell-type hierarchy consisting of 5 levels. When matching the cell types, we used the latent space generated in their original paper (downloaded from https://zenodo.org/record/6337966#.YqmGIidBx3g). When updating the hierarchy with the IPF data, we removed the cell types smaller than 10 cells. Marker genes were downloaded from the lung reference v2 from Azimuth (2, 4). Marker genes for the Meyer cell populations were obtained from [26].

We annotated the fibrosis-specific cell types in greater detail by sub clustering the cell types of interest (macrophages, epithelial cells, myofibroblasts and identifying the subtypes by marker gene expression. We identified transitioning/basaloid epithelial cells by KRT5/KRT17 expression, inflammatory monocyte-derived macrophages by SPP1 expression, and myofibroblasts by the expression of CTHRC1.

## Comparisons

### FR-Match

We ran FR-Match (v2.0.0) with default settings on all pairwise combinations of the PBMC reference datasets (23, 24). Before running FR-Match marker genes have to be selected for each cell type. We do this using the method recommended by the authors of FR-Match: NS-Forest (25). We ran NS-Forest (v3.0) on each dataset separately using the default settings.

### MetaNeighbor

We ran MetaNeighbor (v1.13.0) using the default settings on all pairwise combinations of the PBMC datasets (26). MetaNeighbor returns an AUROC score for all cell-type combinations. As recommended in the MetaNeighbor vignette, we consider two cell types a match when the AUROC is higher than 0.9.

### Azimuth

We run Azimuth using Seurat v4.3.0 (4) and follow the ‘integration_mapping’ vignette.

## Results

### treeArches accurately learns PBMC hierarchy

We showcase treeArches with a simulation where we build a cell-type hierarchy using one bone marrow and three PBMC datasets (19–21, 27) (Table S1). We consider three datasets as the reference (Freytag, Oetjen, and Sun), and one as the query (10X). The annotations of these datasets have been manually harmonized by Luecken et al. (16), so we relabel some cells to enforce the datasets to be annotated at different resolutions (Tabel S2, S3). In the Oetjen dataset, for instance, we relabel all the CD4+ and CD8+ T cells as T cells. The challenge here is to correctly match cell types present in multiple datasets and to reconstruct their hierarchy. Some cell types, however, are dataset-specific and these should thus be added as a new node in the tree. Here, it is important to note that these new cell types are not forced to be aligned with other existing cell types during the integration step and that the classifier used by scHPL contains a good rejection option during the matching step. This harmonizing and afterward relabeling of the cells allows us to manually construct a ground truth hierarchy that we can use to evaluate treeArches (Fig S1).

We remove the batch effects from the reference datasets using scVI (14) and match the cell types in the learned latent space (see **Methods**) (Fig 1C-D, Fig S2). Since both scArches and scHPL are invariant to a different order of the datasets, treeArches will also be invariant (10, 12). For scHPL, however, the datasets still have to be added progressively, which we will do from low to high resolution (Sun - Oetjen - Freytag). The constructed tree by treeArches largely matches the ground truth: seven out of eight Oetjen cell types and all nine Freytag cell types are correctly matched to the Sun cell types (e.g. the CD4+ T cells are a subpopulation of the T cells which are a subpopulation of the lymphoid cells). The six cell types only found in one dataset are all added as new cell types to the tree (e.g. the CD10+ B cells and erythrocytes).

However, the megakaryocyte (MK) progenitor cells from the Freytag and Sun dataset do not match the cells from Oetjen. The Freytag and Sun datasets are PBMC datasets and the Oetjen dataset is a bone marrow dataset. Looking at the expression of marker genes and the location of the megakaryocyte progenitor cells in the UMAP embedding supports our claim that the cell types from Sun and Freytag should not match Oetjen in the hierarchy (Fig S3). Based on marker gene expression, the MK progenitor cells in the Oetjen dataset should be relabeled as early erythrocytes and the MK progenitor cells in the Freytag and Sun dataset as platelets.

After constructing the reference tree from the three datasets, we align the query dataset to the latent space of the reference datasets using scArches and update the learned hierarchy with the new cell types (Fig 1E-F). For this step, only the trained model and reference latent space are needed. Again, almost all cell types (10 out of 12) are added to the correct node in the tree, while the plasma cells and the MK progenitors are added to the tree as new cell types. These cell types contain 21 and 18 cells, respectively, which makes them difficult to match compared to the other cell types in the query dataset, which contain more than 1000 cells on average.

Since there is no method with exactly the same functionality as treeArches, we benchmark parts of the algorithm separately. First, we compare our constructed hierarchy for the reference data to the output of two cell-type matching algorithms: FR-Match and MetaNeighbor (23, 24, 26). It is important to note that these methods were developed for pairwise comparisons and do not construct a hierarchy. We ran both methods on all combinations of the reference datasets and visualized their matches in a graph (Fig S4). To allow comparisons, we transform the learned hierarchy by treeArches to a graph by adding edges between a parent and all descendants (Fig S4). When comparing the resulting graphs to the ground-truth graph constructed based on the relabeled cell types, treeArches outperforms FR-Match and MetaNeighbor (Table S4). Using treeArches, only two edges are missing and no wrong edges were introduced, while using FR-Match and MetaNeighbor there are respectively 11 and 8 wrong edges, and 7 and 11 missing edges.

Next, we compare the cell type classification performance of treeArches to Azimuth (4). Azimuth allows label transfer by projecting a query dataset onto a reference atlas but assumes that the labels of the reference are already harmonized. Therefore, we compare the performance in two ways: 1) using the datasets annotated at a different resolution, and 2) using the datasets with the manually harmonized labels. We use the Sun, Oetjen, and Freytag datasets as a reference and the 10X dataset as the query. In the first comparison, treeArches outperforms Azimuth (Fig S5), but during the second comparison, Azimuth performs better (Fig S6). Here, we also compare the performance of treeArches using the kNN (default) and using a linear SVM which is the best-performing method according to our classification benchmark (28). Since the latent space is not linearly separable anymore, the kNN outperforms the linear SVM (Fig S6). This motivates the use of a kNN classifier within treeArches.

### Increasing the resolution of the human lung cell atlas using treeArches

The human lung cell atlas (HLCA) is a carefully constructed reference atlas for the human respiratory system (2). Sikkema et al. integrated 14 datasets, re-annotated the cells, and constructed a cell-type hierarchy consisting of 5 levels (Fig 2A, Fig S7). Furthermore, they used scArches to project multiple datasets to this reference atlas. Since the cell-type hierarchy for the reference is well-defined, we can omit the reference-building step and leverage treeArches to update the reference hierarchy using one of the labeled query datasets (Meyer) (29). Using scHPL, we matched the cell types of the Meyer dataset to the cell types from the reference (Fig S8). In the updated hierarchy, many cell types from the query dataset match a cell type from the reference as expected based on the cell-type names. Neuroendocrine-Meyer, for instance, is a perfect match to the neuroendocrine from the reference. Since no ground truth cell-type matches between the reference datasets and Meyer is known here, we cannot assess this quantitatively. For some parts of the hierarchy, we can even increase the resolution. If we zoom in on the blood vessel branch in the tree, for instance, the pulmonary and systemic endothelial vascular arterial cell types from the query both match endothelial cells arterial (EC arterial) from the reference (Fig 2B).

**Figure 2.**
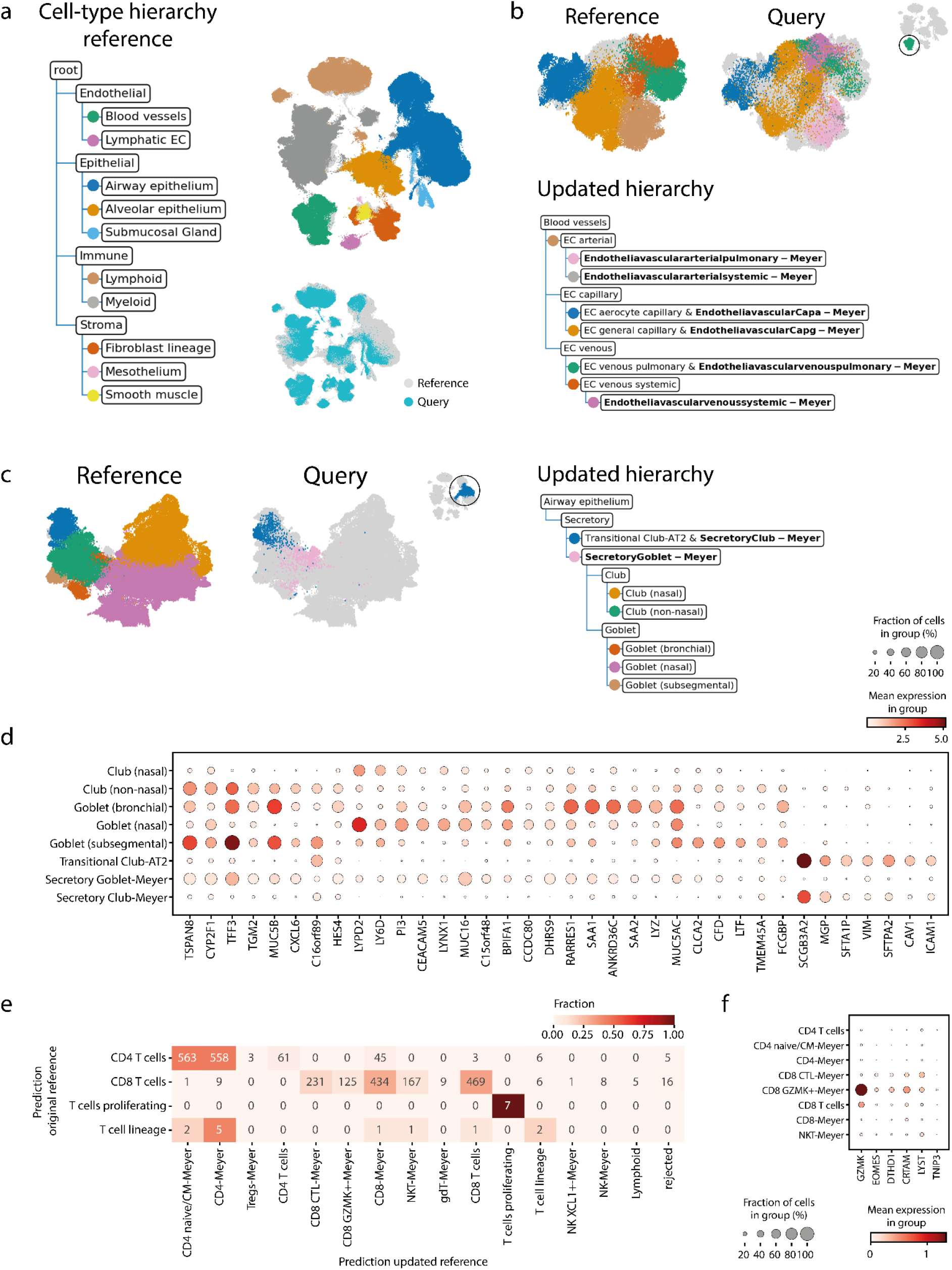
Updated hierarchy when adding Meyer to the reference atlas. **(a)** cell-type hierarchy corresponding to the reference atlas (only the first two levels are shown). Each node represents a cell type in the reference atlas instead of a cell type in a separate dataset of the reference atlas. The UMAP embedding shows the aligned reference and query dataset. The cells in the reference dataset are colored according to their level 2 annotation. **(b-c)** Updated hierarchy zoomed in on the blood vessels and airway epithelium secretory cells respectively. The UMAP embeddings are colored according to their finest resolution. **(d)** Expression of marker genes for club and goblet cells in the reference and query cell types. **(e)** Comparison of the predictions using the original and updated reference on the T-cells of the Tata dataset. **(f)** Expression of marker genes for CD8+ GZMK+ cells.

For some parts of the tree, e.g. the airway epithelium secretory cells, the matches are not what we would expect based on the names (Fig 2C). The secretory goblet cells from the query dataset match not only the goblet but also the club cells from the reference and the secretory club cells match the transitional club-alveolar type 2 (AT2) cells. Transitional club-AT2 cells were only recently discovered, which could explain why they are missing from the original Meyer annotations (30–32). Based on the expression of marker genes, we can conclude that the match between the transitional club-AT2 and secretory club cells is a correct match (Fig 2D). The expression of the marker genes in the other cell types, however, is ambiguous and it is hard to determine what is the correct match. Furthermore, in the HLCA paper, label transfer for these cell types from the reference atlas to the Meyer data did not match well with the original labels either (2).

Furthermore, we see that there are sixteen cell types from the query added to the root node of the tree as a new cell type (Fig S8). Of these cell types, most of them, e.g. chondrocytes, erythrocytes, Schwann cells, and B plasmablasts, are indeed not in the reference atlas. For some, such as some macrophage subtypes that are seen as new, it is more difficult to determine whether they are new or whether they should match one of the macrophage subtypes in the tree. The ‘Macro CHIT1’ cells from the Meyer dataset, for instance, form a relatively big cell type of 1570 cells and are still seen as new. We visualized the expression of *CHIT1*, the gene this cell type was named after, and the marker genes that were used to annotate the cells in the reference data (Fig S9). This shows that the Macro CHIT1 cell type is the only cell type that expresses *CHIT1*. Furthermore, the marker gene profile of the other cell types does not correspond to the profile of the Macro CHIT1 cells, which indicates that this cell type was indeed rejected correctly.

However, twelve out of 77 cell types are missing from the tree, which means that it was impossible to match these Meyer cell types with a cell type from the reference. Due to many-to-many matches between the reference and query cell types, it is sometimes unclear where a cell type should be added to the tree. Especially, when the boundary between cell types is diffuse, it can be quite arbitrary where to put the threshold. If this threshold is different in each dataset or if cells are wrongly annotated in general, this can cause impossible matching scenarios. Here, we notice that this mainly happens with some immune and stromal subtypes. The B cells and plasma cells from the reference and Meyer dataset, for instance, could not be matched automatically, which is caused by the plasma cells in the Meyer dataset that are partially misannotated (Fig S10).

Next, we annotate a second healthy query dataset (Tata) (31) using the original and updated reference to show that cells in this new query dataset will indeed be mapped to the new Meyer cell types we added to the hierarchy. The majority of the predictions remained unchanged (72.1%, Fig S11). When the predictions differ, cells are often annotated as a Meyer cell type which is a subpopulation of the original annotation (18.4%). A clear example is the T cells: cells previously annotated as CD4+ or CD8+ T cells are now annotated as a subpopulation (Fig 2e). These new annotations are supported by the expression of marker genes (Fig S12, Fig 2f).

### treeArches identifies unseen disease-associated cell types in the query data

Next, we show how we can use treeArches to detect previously unseen cell types in idiopathic pulmonary fibrosis (IPF) samples (33). This dataset was mapped on the HLCA with scArches (Fig 3A-C). Ideally, we would use scHPL to update the hierarchy with the cell types from this query dataset. A downside of the original annotations, however, is that the resolution is very low. Cells are, for instance, only annotated as endothelial cells. Therefore, we used scHPL to predict the labels of the IPF data and compare those predictions to the original annotations (Fig 3D). In the predictions, we see some interesting differences between the IPF and healthy cells.

**Figure 3.**
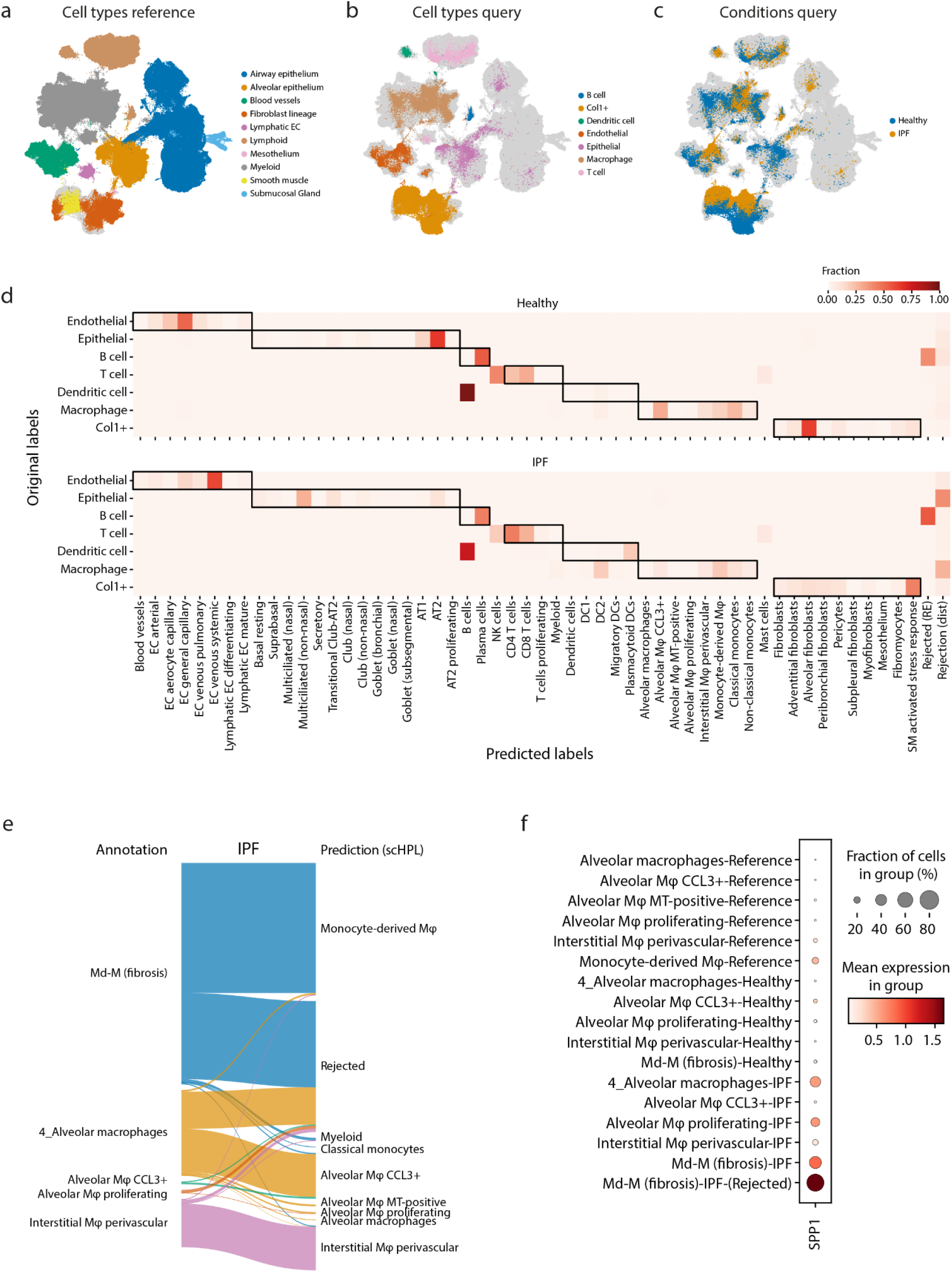
Identifying diseased cells in IPF data. **(a-c)** UMAPs show the HLCA and IPF datasets after alignment. The cells are colored according to their cell type or condition. **(d)** Heatmap showing the predicted labels by scHPL and original labels. The dark boundaries indicate the hierarchy of the reference tree. **(e)** Sankey diagram showing the new annotations and predictions for the macrophages for the IPF condition. **(f)** Expression of *SPP1* in the different cell types of the reference and query datasets.

For the IPF cells, many macrophages and epithelial cells are rejected, while almost none for the healthy cells. Furthermore, most healthy Col1+ cells are predicted to be alveolar fibroblasts, while the diseased Col1+ are mainly SM-activated stress response cells. In all datasets, however, we notice confusion between the B cells and dendritic cells. Based on marker gene expression, the cells originally annotated as B cells and dendritic cells are more likely to be plasmablasts and B cells respectively (Fig S13). The cells originally annotated as dendritic cells also overlap in the UMAP with the lymphoid lineage mainly instead of the myeloid lineage (Fig 3A-B).

Next, we annotated the cells at a higher resolution (see **Methods**) and used these annotations to update the hierarchy (Fig S14). In the updated hierarchy, the healthy and IPF transitioning epithelial cells are not present in the reference atlas and are now correctly added as a new cell type. As expected, we also see some differences in how the healthy and IPF cell types were added to the tree. IPF alveolar macrophage proliferating cells, for instance, are seen as new, while the healthy cells match with the same cell type in the hierarchy. For other IPF macrophage cell types, however, this is not the case even though many cells were rejected previously. Comparing the new annotations with the previously obtained predictions and the matches in the hierarchy, we notice that there are still many macrophages rejected (Fig 3E). For most IPF cell types, however, only a subset of the cells is rejected. For instance, for the IPF monocyte-derived macrophages (Md-M), 486 cells are rejected and 750 are predicted to be Md-M. Therefore, the two cell types are still matched. Comparing the two IPF ‘subtypes’ of Md-M, the top differentially expressed gene is *SPP1*. Monocytes and macrophages expressing *SPP1* are known to be a hallmark of IPF pathogenesis (34, 35). The rejected Md-M cells are the only group of cells expressing *SPP1* (Fig 3F). By combining the confusion matrices with the created hierarchy, these diseased subtypes are easily found, either directly as the proliferating cells, or by looking at the rejected cells of a matched cluster.

### treeArches can correctly map cell types across species

Next, we show how treeArches can be applied to map the relationship between cell types of different species. We construct a cell-type hierarchy for the motor cortex of the brain using human, mouse, and marmoset data (Table S5) (22). We integrate the reference datasets, mouse and marmoset, using scVI and construct the cell-type hierarchy for the GABAergic neurons using scHPL (Fig 4A-B). Here, we only focus on this subset to make the results less cluttered. Almost all cell types (5 out of 7) are a perfect match, except for ‘Meis2’ and ‘Sncg’. In the latent space, the Meis2 cell types from mouse and marmoset also show no overlap, and both cell types were defined using different marker genes (Fig S15A-B). Furthermore, Bakken et al. also didn’t find a match between these two (22). It is less clear why the Sncg cell types (559 and 960 cells in mouse and marmoset respectively) do not match. Even though the cell types are aligned in the UMAP embedding as expected and the marker genes correspond quite well, the cells are rejected based on distance (Fig S15C-D). This means that the cells are still too separated in the latent space. Next, we align the human dataset to the reference using architectural surgery and add the human cell type to the reference hierarchy (Fig 4B-C). Here, the constructed hierarchy looks like what we would expect based on the names of the cell types.

**Figure 4.**
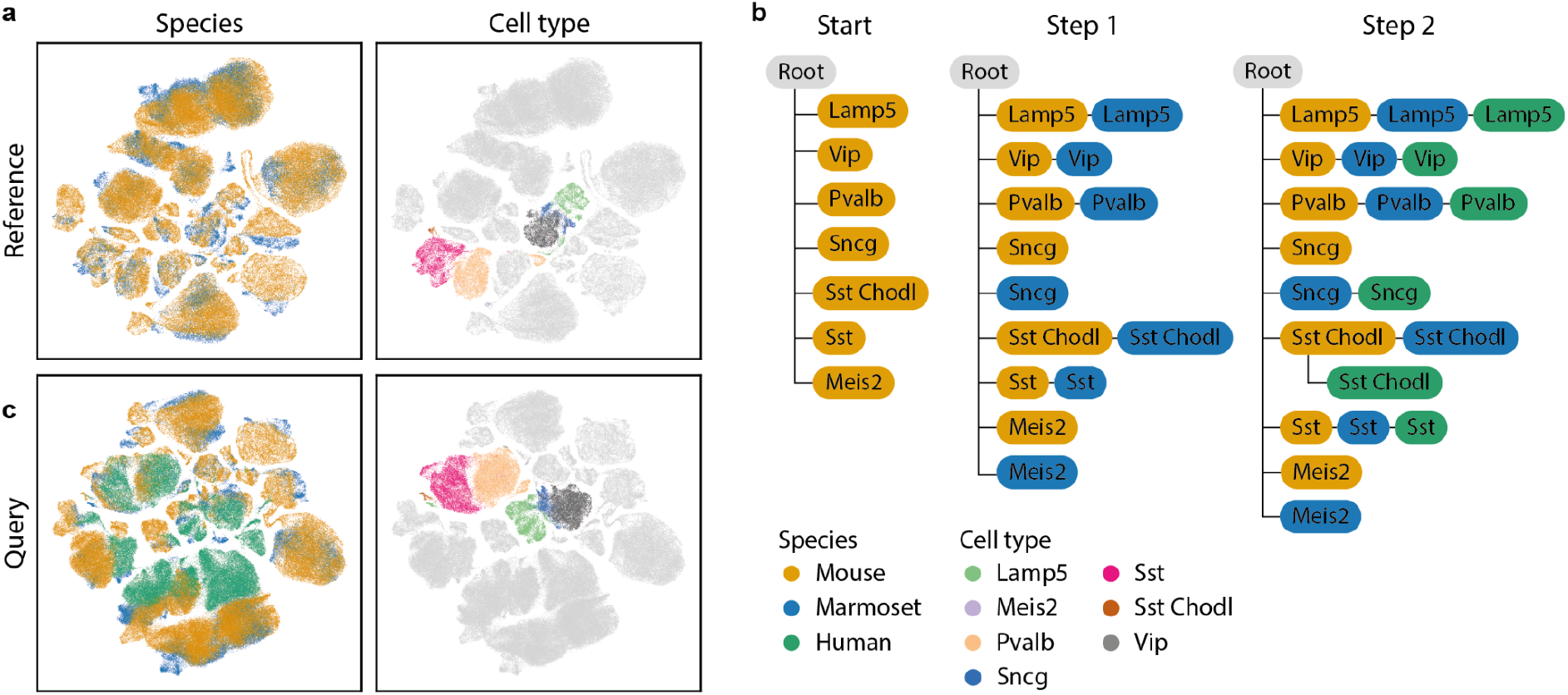
Results motor cortex across species. **(a)** UMAP embedding of the integrated reference datasets. **(b)** Learned hierarchy when combining mouse and marmoset (step 1) and after adding human (step 2). The color of each node represents the dataset(s) that the cell type originates from. **(c)** UMAP embedding after architectural surgery with the human dataset.

All previous results were obtained using the default parameters (number of neighbors = 50, see **Methods**), which turned out to be relatively robust (Fig S16). The main difference is whether a match is found between the Sncg cell types. When increasing the number of neighbors, this match is correctly found.

## Discussion

In this study, we present treeArches, a method to create and extend a reference atlas and the corresponding cell type hierarchies. treeArches builds on scArches, which allows users to easily map new query datasets to the latent space learned from the reference datasets using architectural surgery. Architectural surgery has the advantage that the reference datasets are not needed anymore for the mapping and that the latent space corresponding to the reference datasets does not change. This last point is especially important for scHPL, which then allows users to match the cell types of multiple labeled datasets to build a cell-type hierarchy. If the latent space of all datasets would be altered when a new dataset is added, we would have to restart the construction of the tree completely.

We have shown three different situations where treeArches can be applied: building a reference atlas from scratch, extending an existing reference atlas to add new cell types or increase the resolution, or using an existing reference atlas to label cells in a new dataset. By using the HLCA data, we show an example of how treeArches can be used to extend a hierarchy or to label cells in a new dataset. The HLCA reference atlas consists of 16 datasets with a well-defined cell-type hierarchy. We show that treeArches can be used to extend this hierarchy. For instance, by increasing the resolution of some branches of the tree, but also by adding new cell types. We could also detect diseased cell types in the IPF datasets.

Whether building or extending a reference atlas or labeling new cells, it is essential that we can detect new cell types, such as disease-specific cell types. To do so, it is important that during the mapping, the cell types are not forced to align; the biological variation should be preserved. Furthermore, during the classification, there should be a correctly working rejection option (i.e. cells are recognized to belong to a new unseen class). Here, we showed that this indeed works in all tested scenarios.

Due to the extended rejection options, however, it is difficult to match small cell types (less than 50 cells). We modified the kNN classifier from scHPL such that the number of neighbors automatically decreases when there is a small cell type in the training data, but apparently, this is not sufficient in all cases. The number of neighbors is a trade-off between the ability to learn a representation for small cell types and the generalizability of the big cell types.

treeArches relies on the original annotations to extend the cell-type hierarchy. This can be a problem in two different situations. If the annotations are missing or at a too low resolution, it is impossible to extend the atlas. This was the case with the original annotations of the IPF dataset. Alternatively, annotations can have a high resolution, but (partially) incorrect. Especially when there is no clear boundary between cell types, experts might disagree on where to put the boundary (the threshold for the classifier). Inconsistencies like this might result in a hierarchy that looks erroneous at first sight. In those cases, however, treeArches can still be more useful than expected. A cell-type hierarchy that looks different than expected, is usually a sign that the original annotations are inconsistent (e.g. different thresholds are used in different datasets). Certain parts of the dataset, e.g. the cell types that could not be added to the tree or caused confusion, can then be reannotated. Furthermore, the tree can still be adapted afterwards. Examples of this are the goblet and club cells in the HLCA and the megakaryocyte progenitor cells in the PBMC datasets. The learned hierarchy is a good starting point. Based on marker gene expression or expert knowledge, cell types can also be added to the tree, removed from the tree, or rewired. After manually adapting the tree, the classifiers have to be retrained though.

Our proposed method builds upon existing data integration methods. Thus, it naturally inherits both advantages and disadvantages linked to these existing models. As previously reported (12), the choice of the reference building algorithm and reference atlas itself can influence the quality of reference mapping. Therefore, in scenarios where the query dataset is strikingly different from the reference, the integrated query will still contain batch effects leading to inaccurate estimation of hierarchies in treeArches. This erroneous modeling results in weak label transfer results and thus identifies many overlapping cell types between query and reference as a new cell type only present in the query. We advise users to choose a comprehensive reference atlas and extensively benchmark and screen various data integration methods for an optimal reference representation (16).

In summary, we present treeArches, a method that can be used to combine multiple labeled datasets to create or extend a reference atlas and the corresponding cell-type hierarchy. This way we provide users with an easy-to-use pipeline to map new datasets to a current reference atlas, match cell types across multiple labeled datasets, and consistently label cells in new datasets. With the increasing availability of reference atlases, we envision treeArches facilitating the usage of reference atlases allowing users to automatically analyze their datasets from label transfer to the automatic identification of novel cell states in the query data. In conclusion, treeArches will enable a data-driven path towards consensus-based cell type annotation of (human) tissues and will significantly speed up the building and annotation of atlases.

## Code availability

treeArches is part of the scArches repository (https://github.com/theislab/scarches). The code for scHPL as a standalone package can be found here: https://github.com/lcmmichielsen/scHPL. All code to reproduce the results and figures can be found at the reproducibility GitHub: https://github.com/lcmmichielsen/treeArches-reproducibility

## Data availability

PBMC count data: https://drive.google.com/uc?id=1Vh6RpYkusbGIZQC8GMFe3OKVDk5PWEpC

Brain count data: https://doi.org/10.5281/zenodo.6786357

PBMC + brain latent space: https://doi.org/10.5281/zenodo.6786357

HLCA latent space: https://zenodo.org/record/6337966#.YqmGIidBx3g

## Acknowledgments

ML acknowledge useful feedback from Malte Luecken regarding the experiments. We also appreciate Sergey Rebakov’s contributions in merging the code within the scArches code base.

## Author contributions

ML conceived the project with contributions from LM and AM. ML, LM, AM designed the experiments. LM performed the experiments. DS helped with the analysis of the IPF data. LS provided feedback essential to HLCA analysis. LM and ML wrote the manuscript. AM, MJTR, FJT supervised and approved the manuscript.

## Competing interest

F.J.T. consults for Immunai Inc., Singularity Bio B.V., CytoReason Ltd, and Omniscope Ltd, and has ownership interest in Dermagnostix GmbH and Cellarity.

## Funding

This research was supported by an NWO Gravitation project: BRAINSCAPES: A Roadmap from Neurogenetics to Neurobiology (NWO: 024.004.012) and by grant number 2019-002438 from the Chan Zuckerberg Foundation, by the European Union’s Horizon 2020 research and innovation programme under grant agreement No 874656, by the Helmholtz Association’s Initiative and Networking Fund through Helmholtz AI [ZT-I-PF-5-01] and by the Bavarian Ministry of Science and the Arts in the framework of the Bavarian Research Association “ForInter” (Interaction of human brain cells).

## Supplementary Material

**Figure S1:**
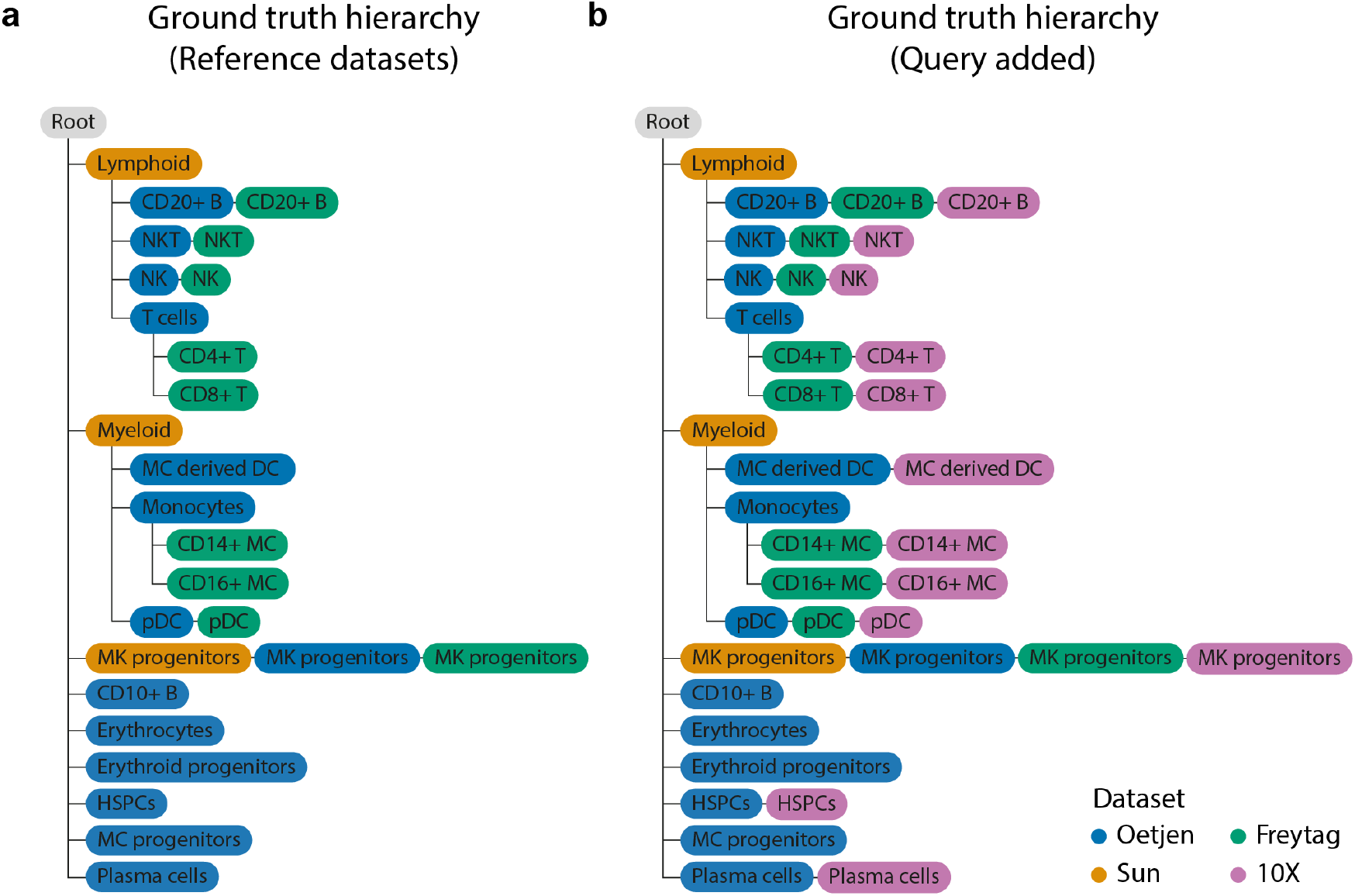
Ground truth hierarchy for a) the reference datasets (Sun, Oetjen, and Freytag) and b) when adding the query dataset (10X).

**Figure S2:**
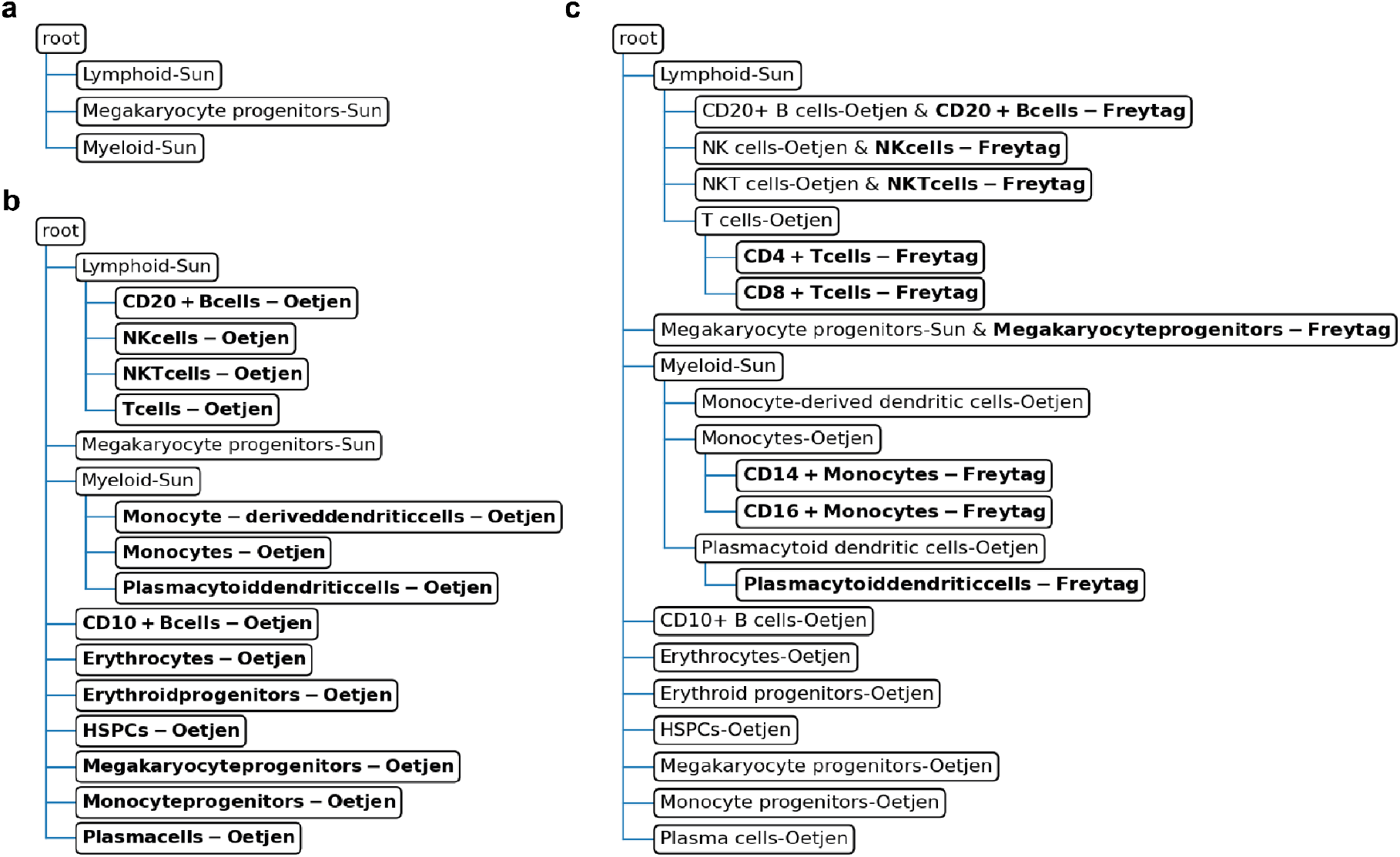
Intermediate step when creating the cell-type hierarchy for the reference PBMC datasets. a) Starting tree, which is a flat tree containing only the cell types of the Freytag dataset, b) Oetjen dataset added, c) Sun dataset added

**Figure S3:**
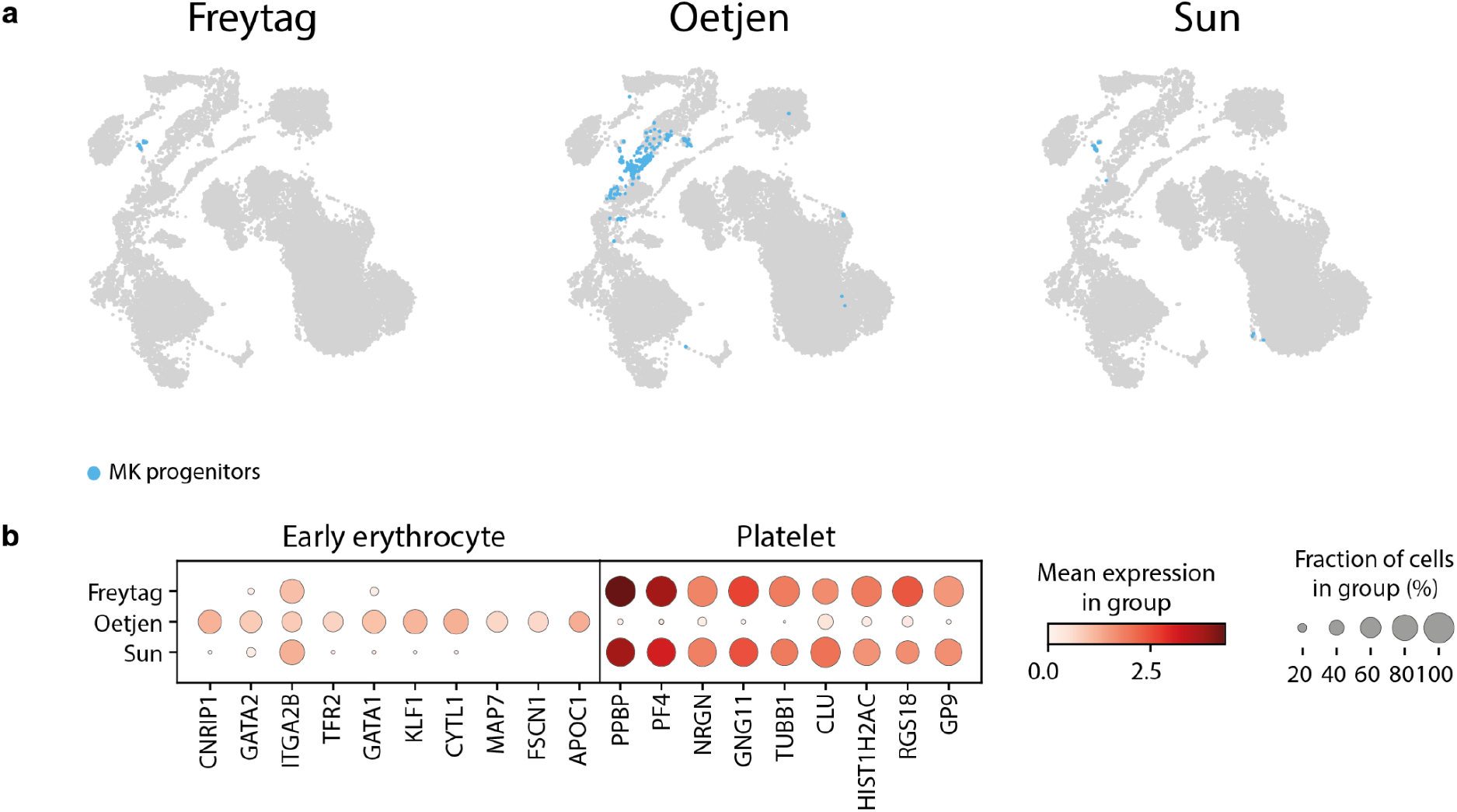
a) UMAP embedding showing the different cell types in the Freytag, Oetjen, and Sun dataset. Megakaryocyte (MK) progenitor cells of the Freytag and Sun dataset are at a different location than the Oetjen dataset. b) Marker gene expression for early erythrocytes and platelets in the three datasets.

**Figure S4.**
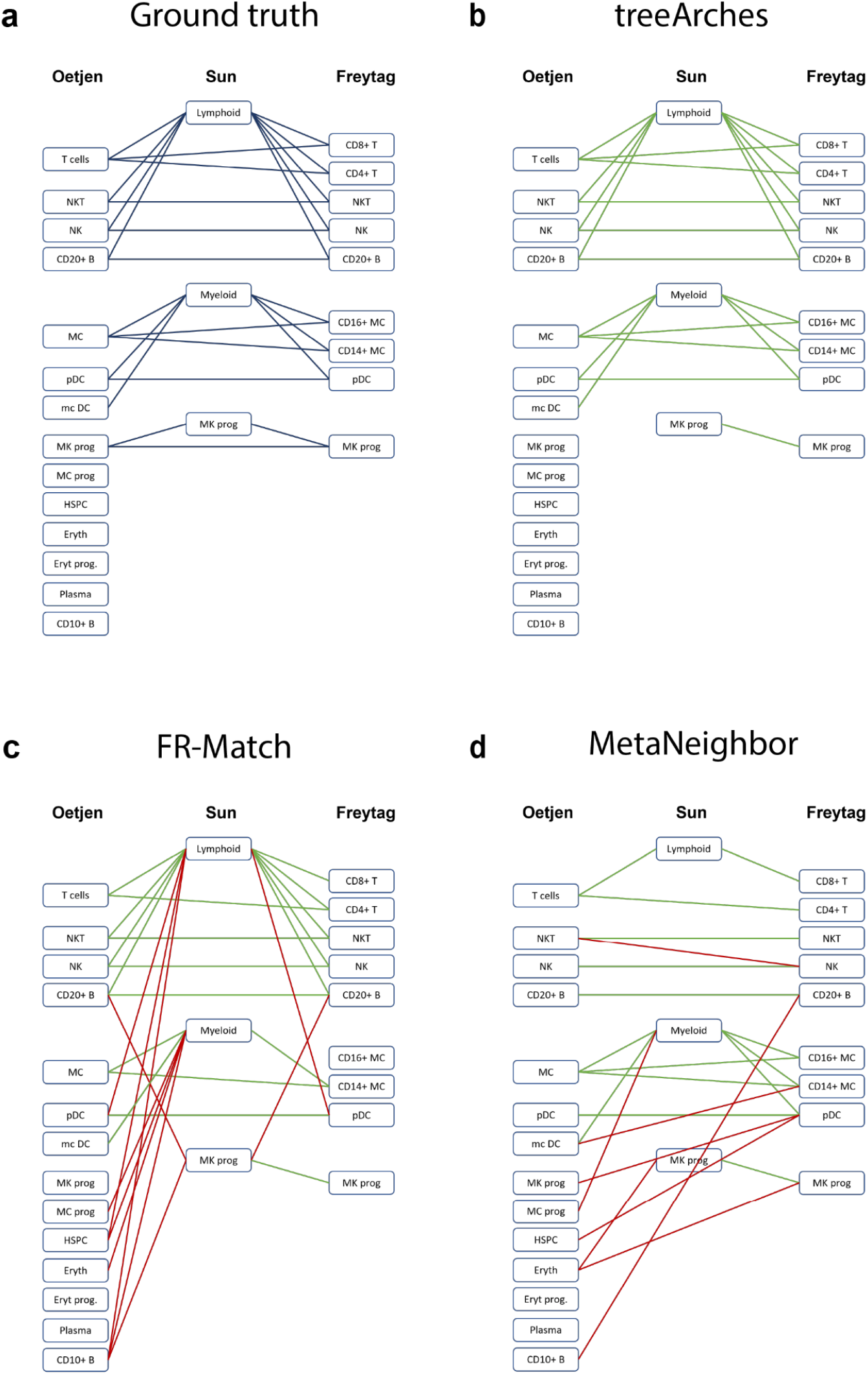
Comparison of cell type matching algorithms. a) Graph showing the ground truth matches between cell types. b-d) Results of treeArches, FR-Match, and MetaNeighbor respectively. Edges in green indicate a right edge, and edges in red indicate a wrong edge.

**Figure S5:**
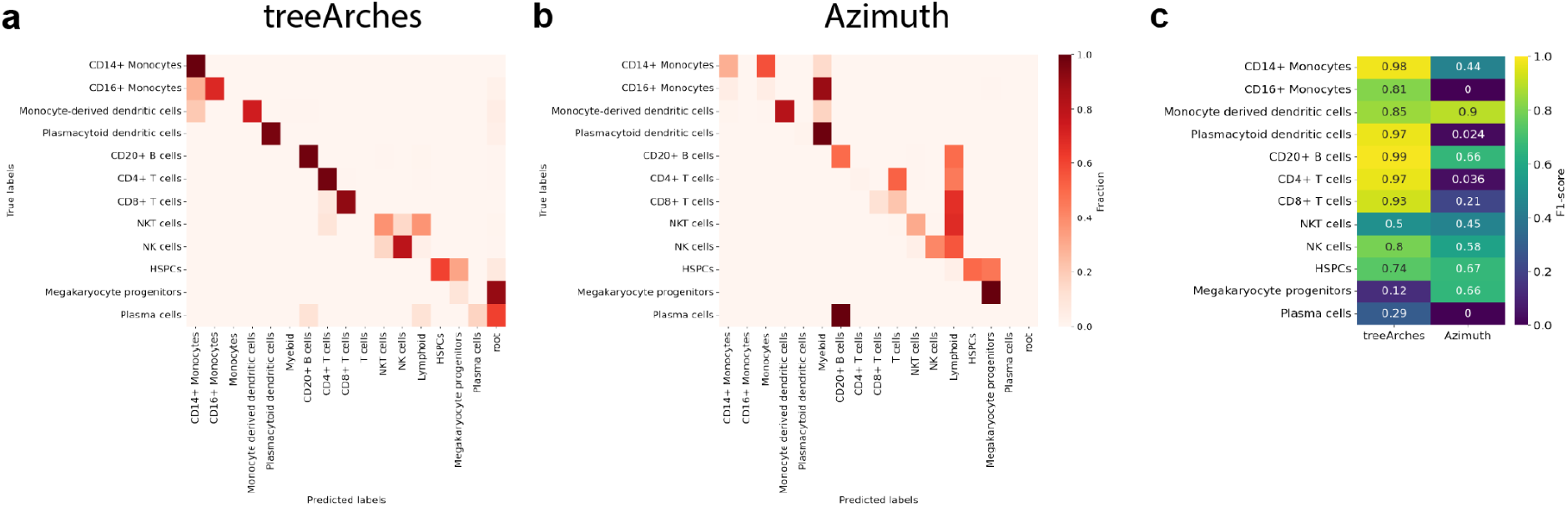
Comparison of classification performance using hierarchical labels. a-b) Confusion matrix for treeArches and Azimuth. c) F1 scores per cell type.

**Figure S6:**
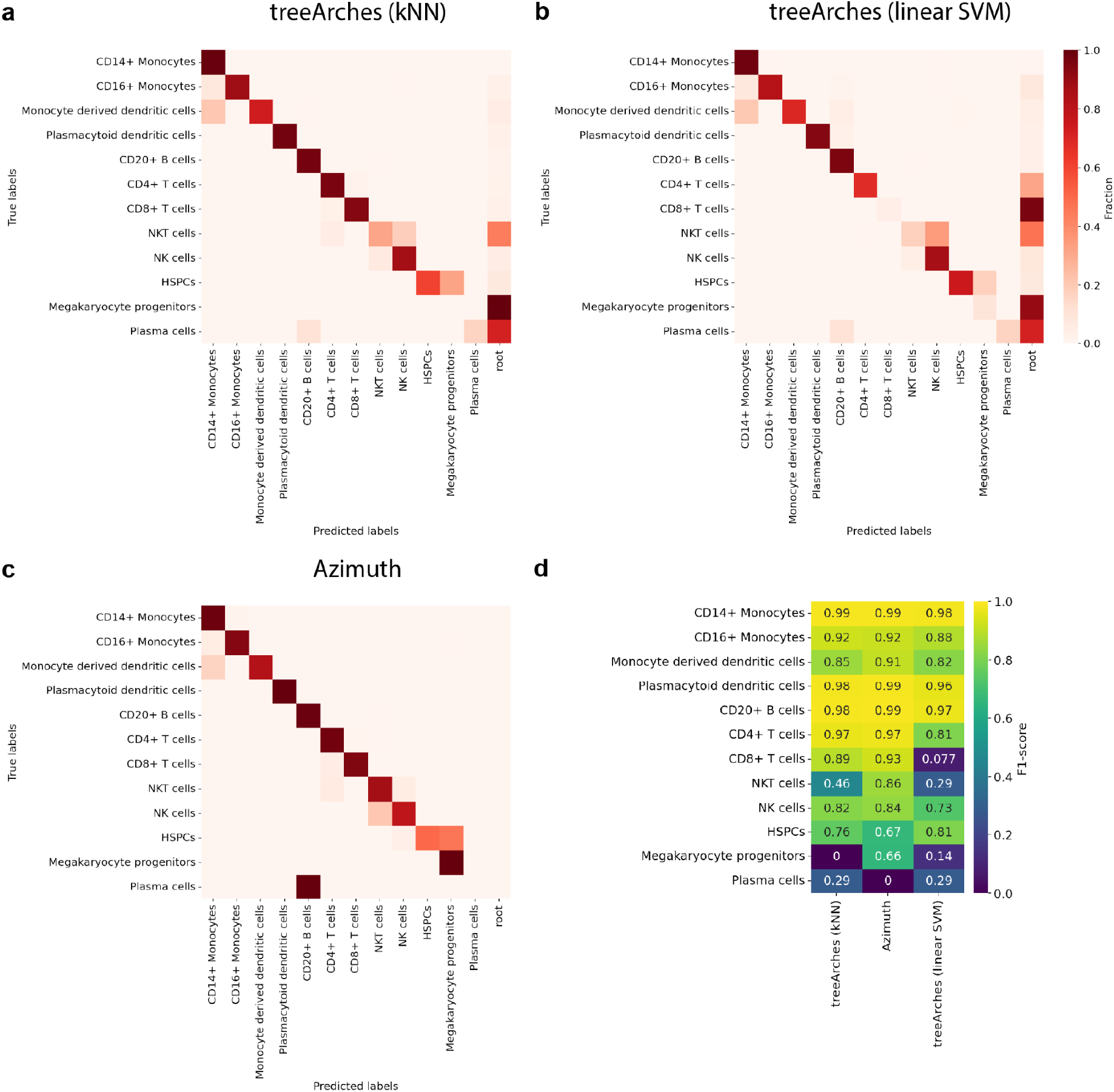
Comparison of classification performance using the harmonized labels. a-c) Confusion matrix for treeArches with kNN, treeArches with linear SVM, and Azimuth. d) F1 scores per cell type.

**Figure S7:**
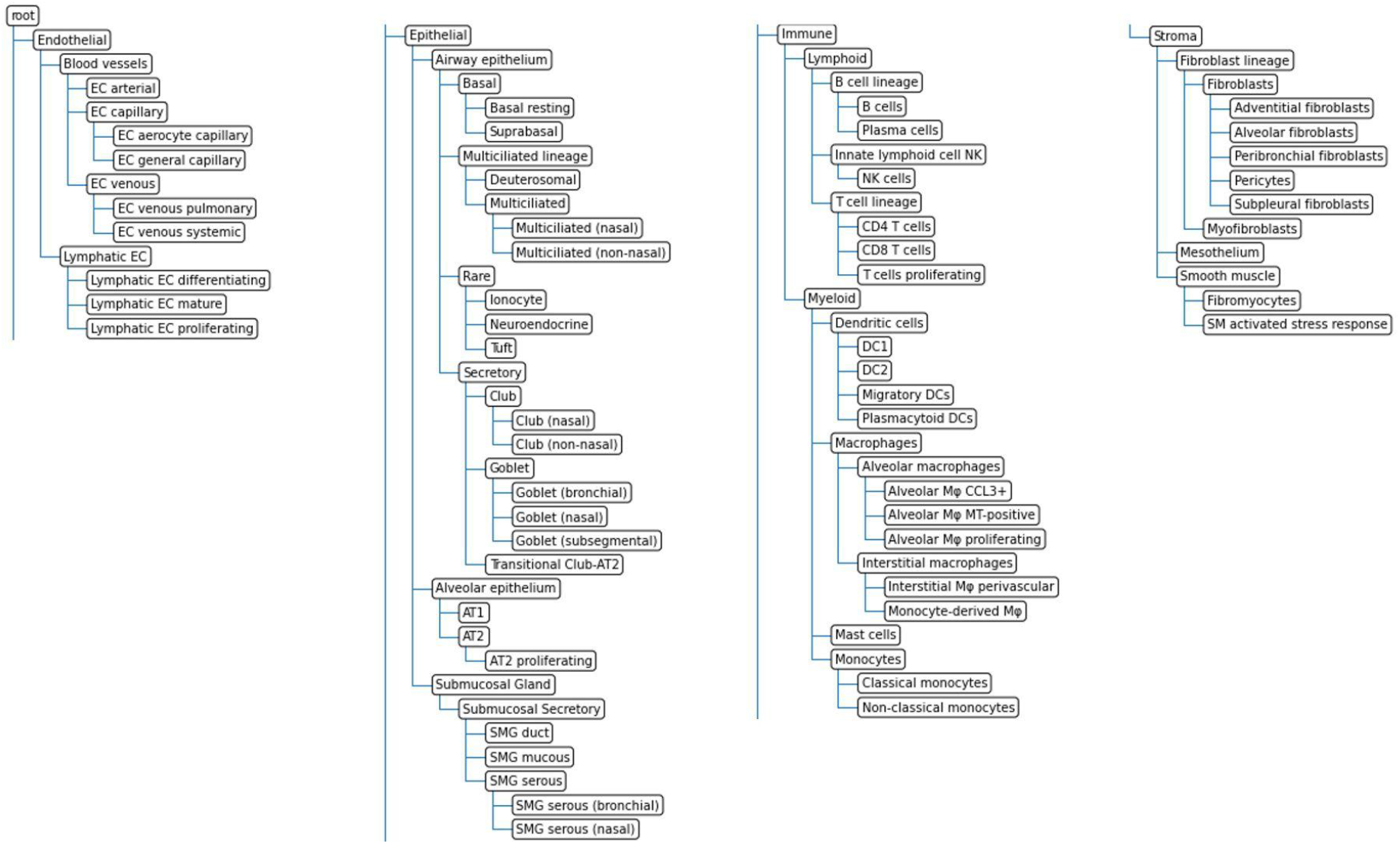
cell-type hierarchy constructed for the reference atlas (2).

**Figure S8.**
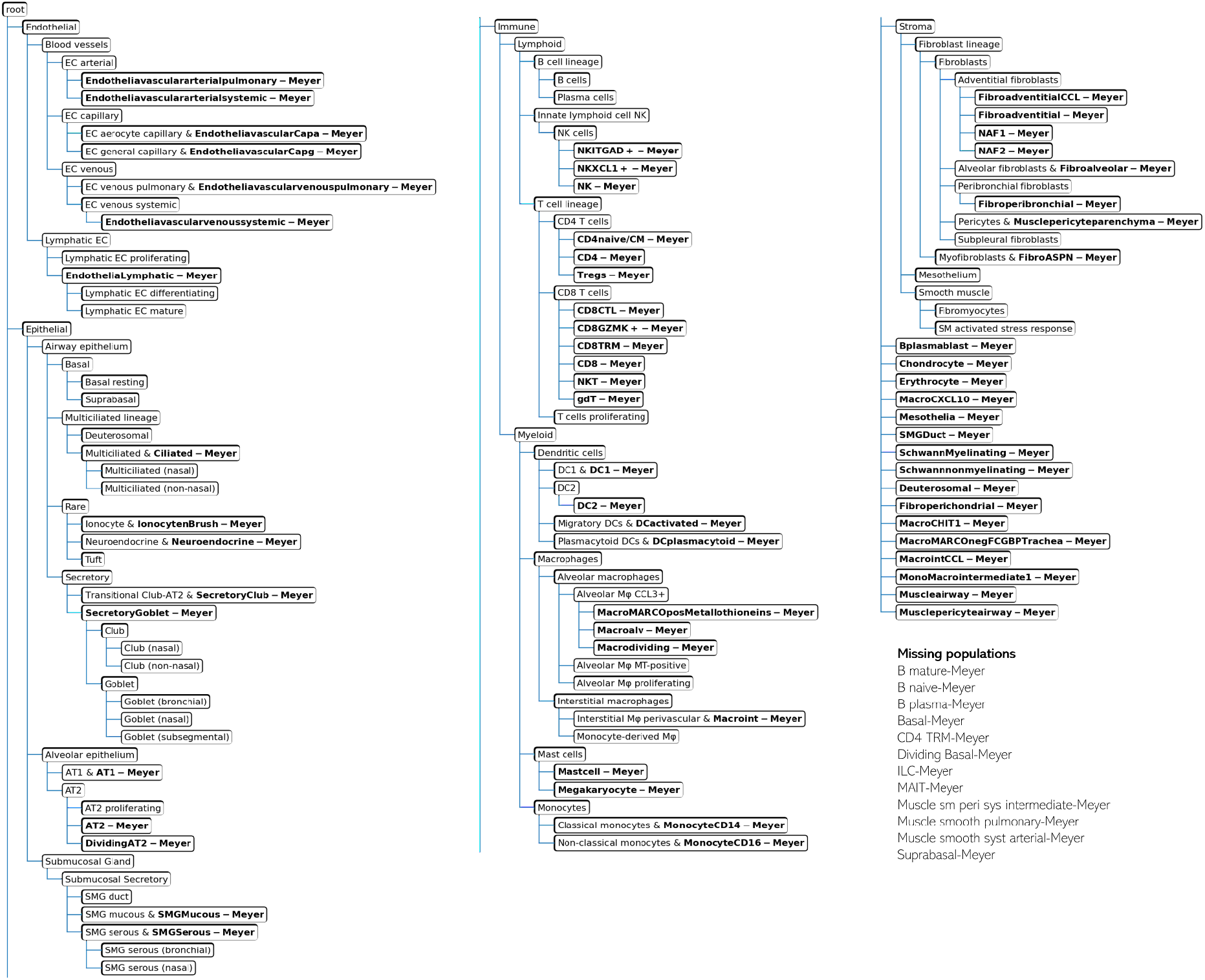
Updated cell-type hierarchy learned by adding a query dataset (Meyer dataset) to the reference tree.

**Figure S9.**
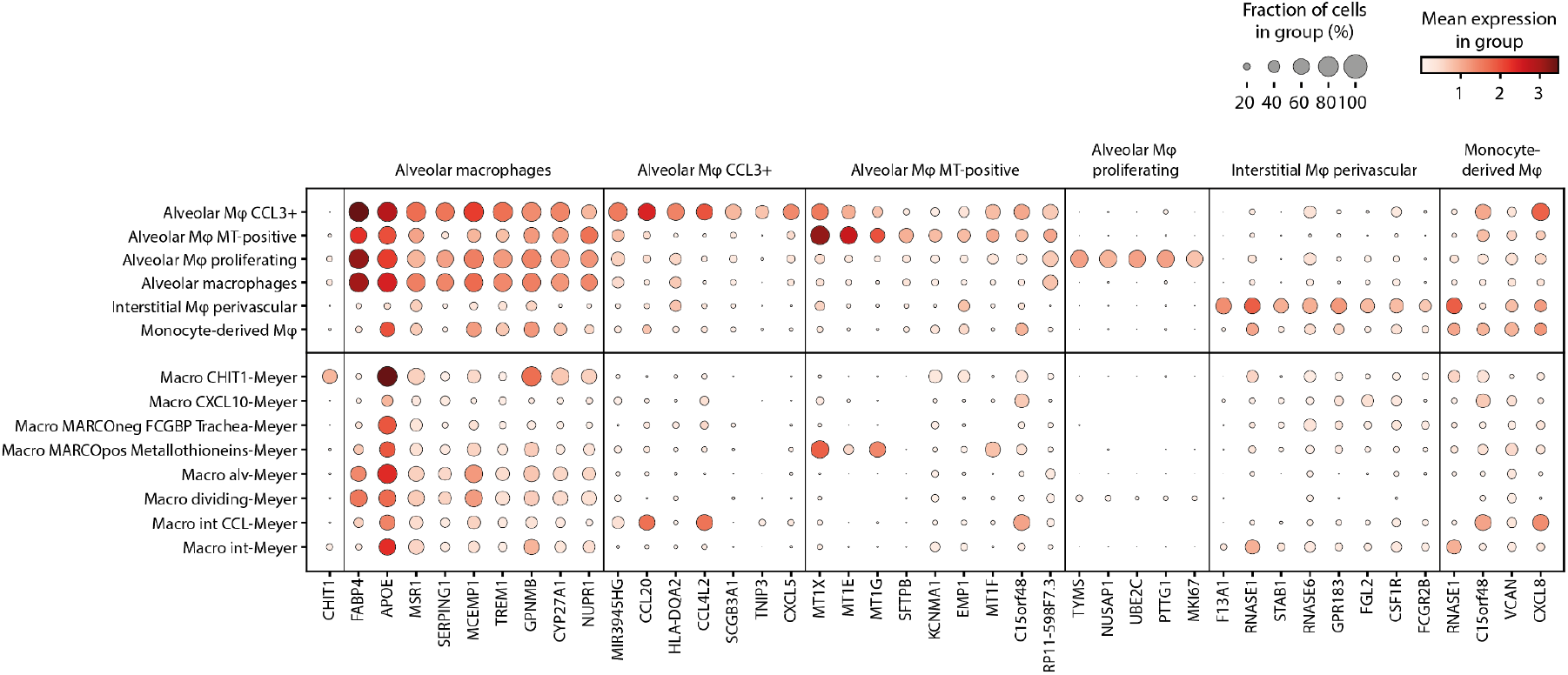
Marker gene expression for macrophage cell types in the reference datasets and Meyer dataset. The first column shows the expression of *CHIT1*, a gene used to annotate the Macro CHIT1 cells in the Meyer dataset. The rest of the genes are grouped according to the cell type in the reference atlas they were used as a marker for.

**Figure S10.**
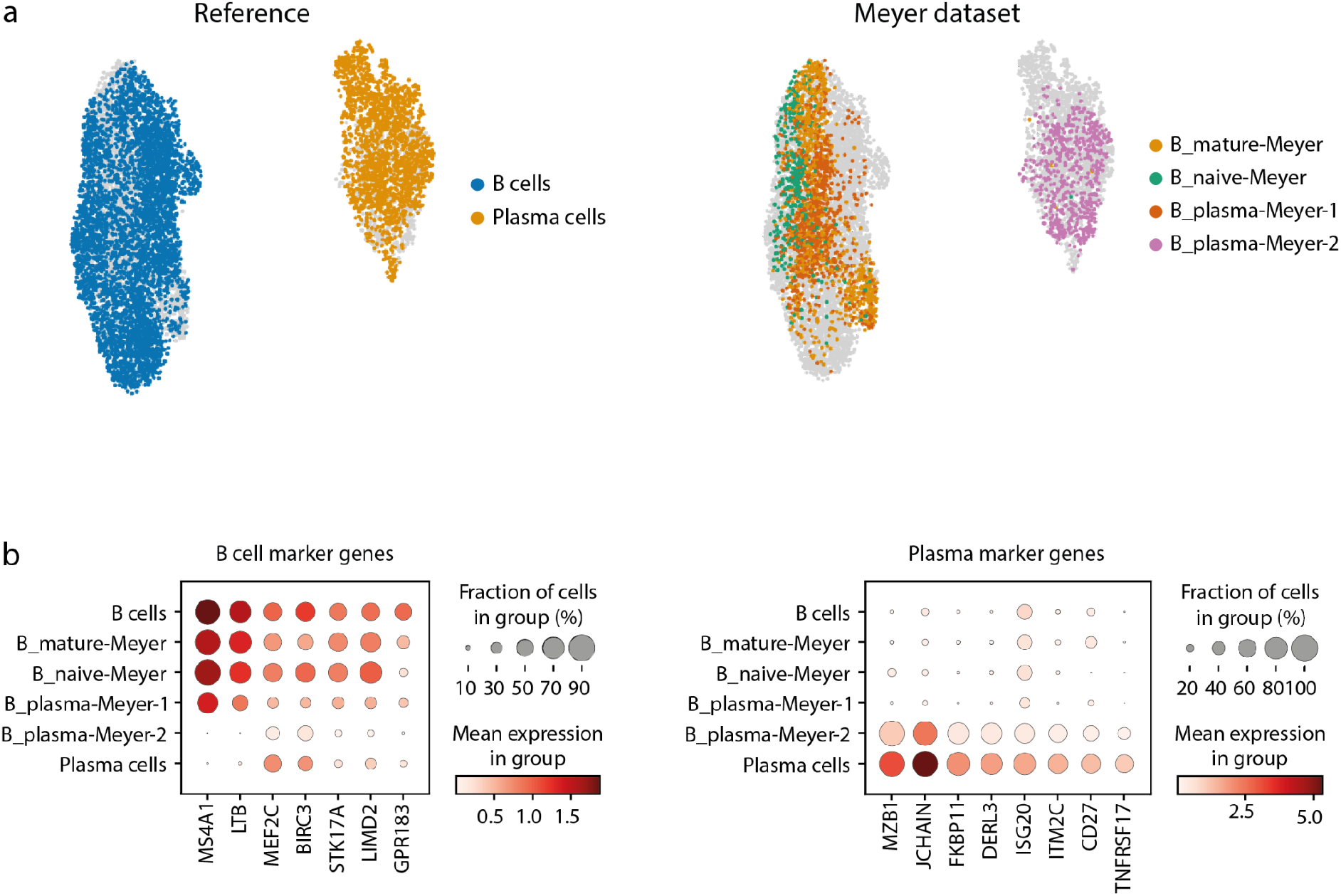
a) UMAPs showing the B cells and plasma cells in the reference and Meyer dataset. We split the plasma cells in the Meyer dataset into two groups. The first group overlaps with the reference B cells and the second group overlaps with the reference plasma cells. b) B cell and plasma cell marker gene expression in the reference and Meyer cell types. Plasma-1 from Meyer shows B cell marker gene expression, while Plasma-2 from Meyer shows plasma marker gene expression.

**Figure S11.**
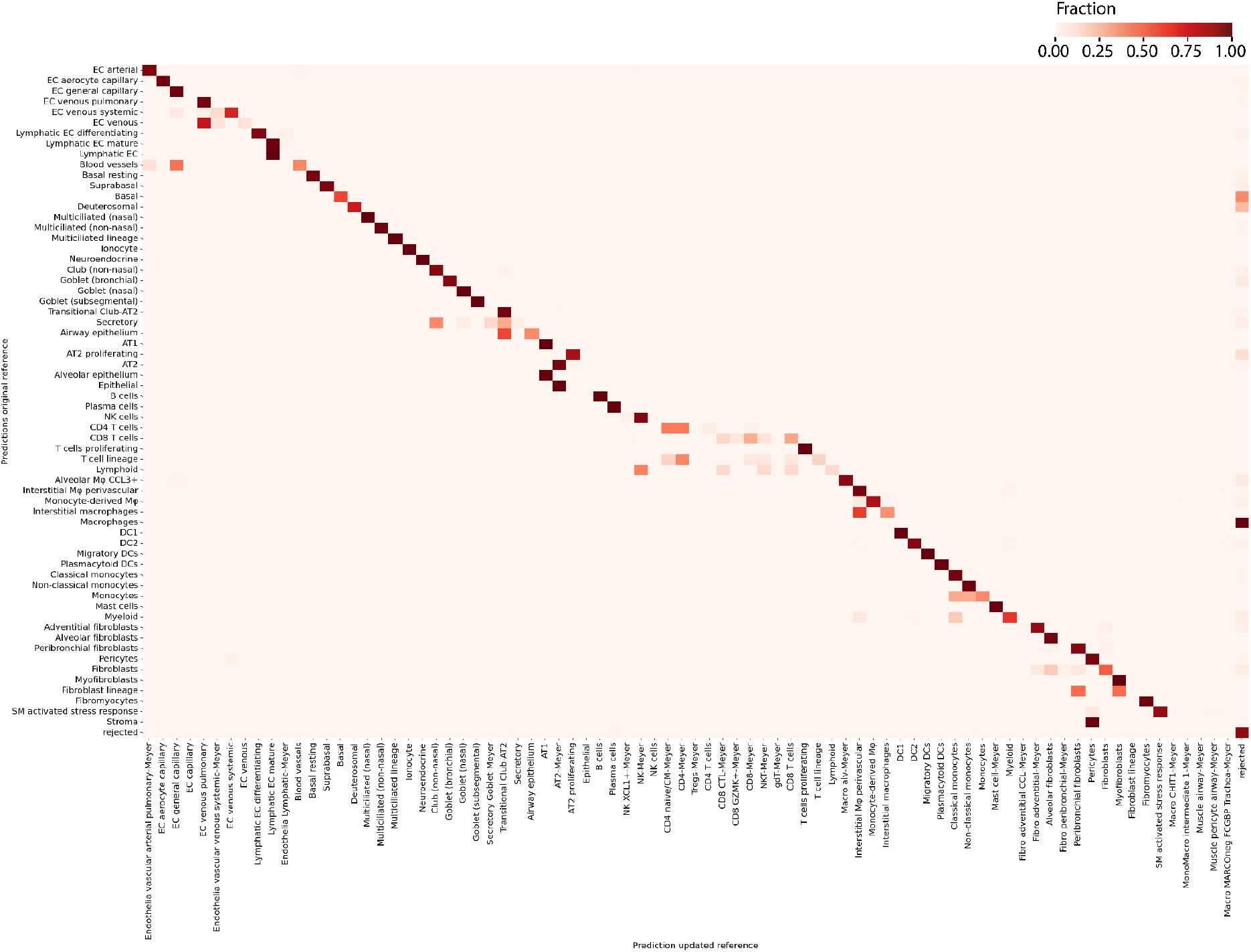
Confusion matrix comparing the predictions on the Tata dataset using the original reference and the updated reference.

**Figure S12.**
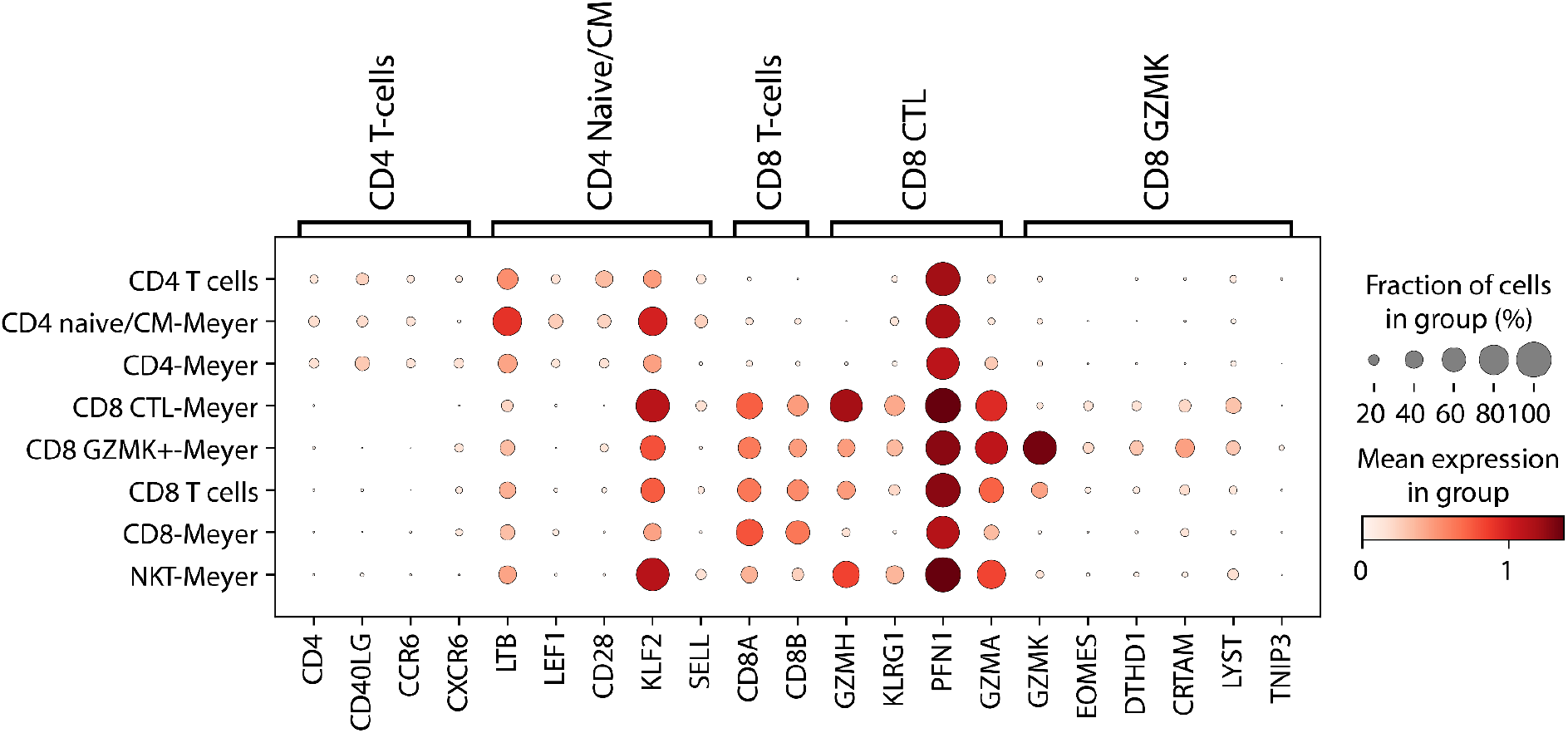
Expression of marker genes for CD4+ T cells, CD4+ naive/CM, CD8+ T cells, CD8+ CTL and CD8+ GZMK.

**Figure S13.**
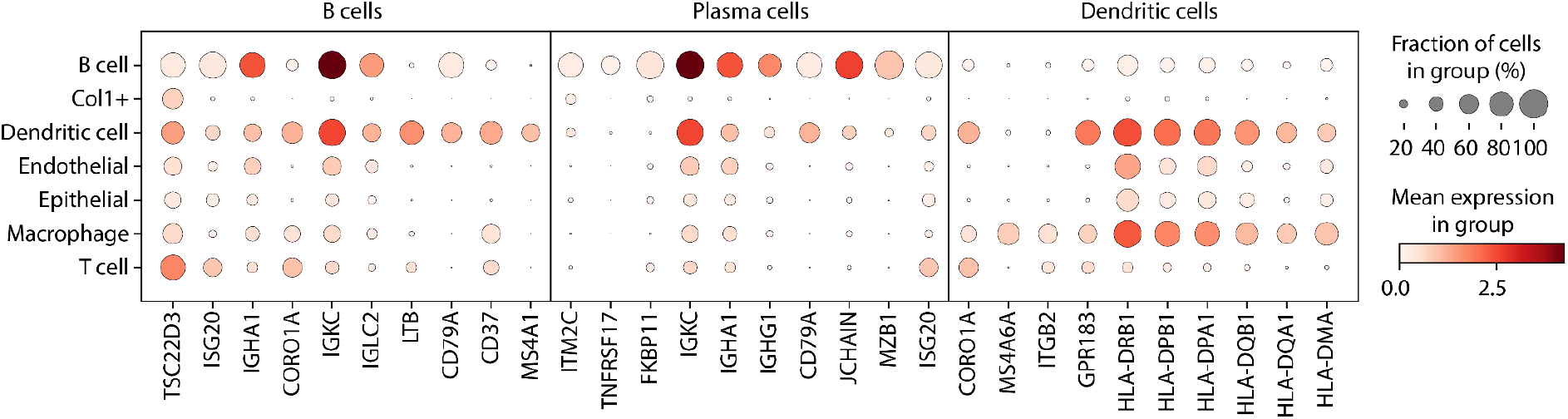
Expression of marker genes for B cells, plasma cells, and dendritic cells in the cell types in the IPF dataset.

**Figure S14.**
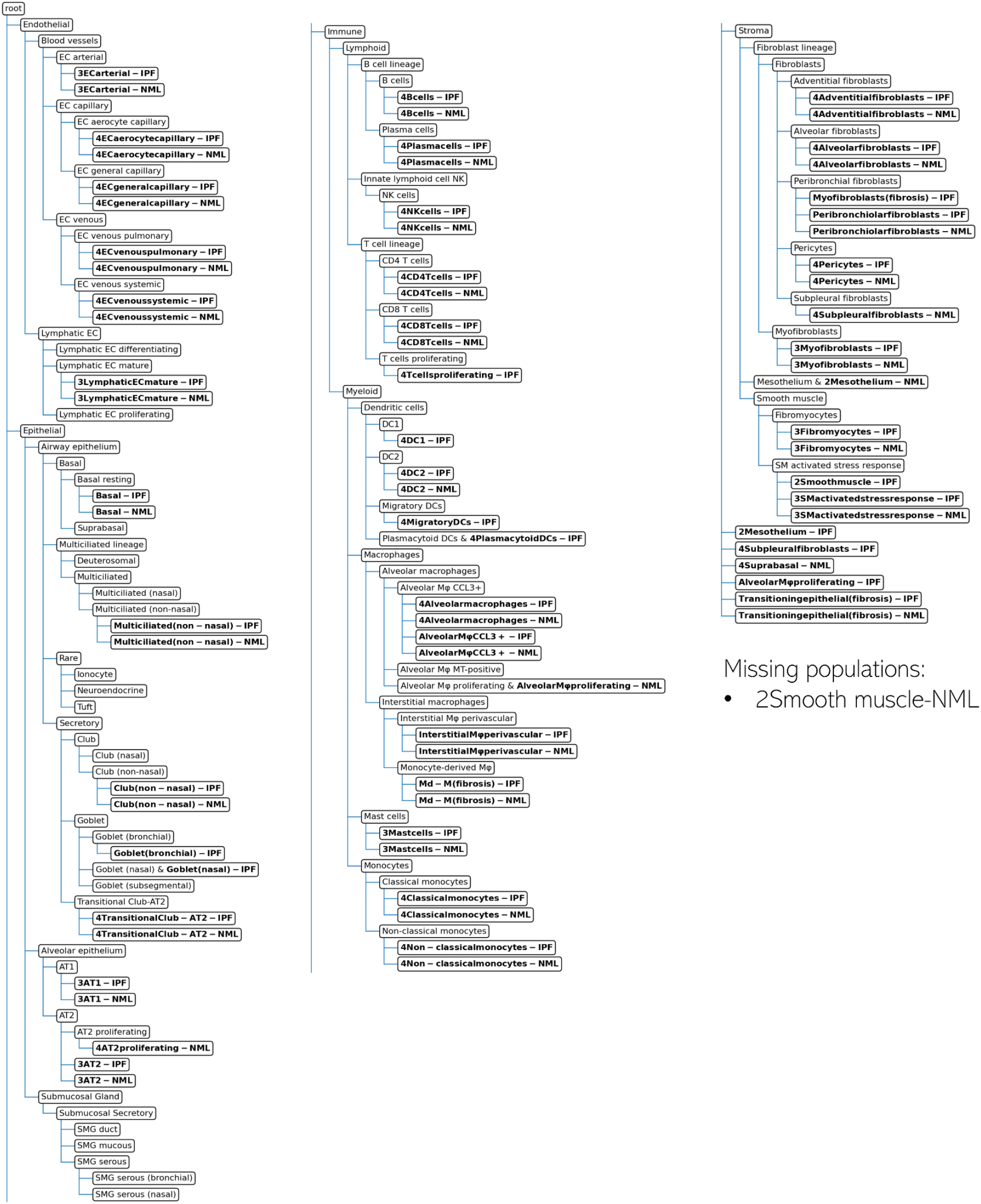
Updated hierarchy of the HLCA after adding the IPF dataset (IPF condition and normal condition).

**Figure S15:**
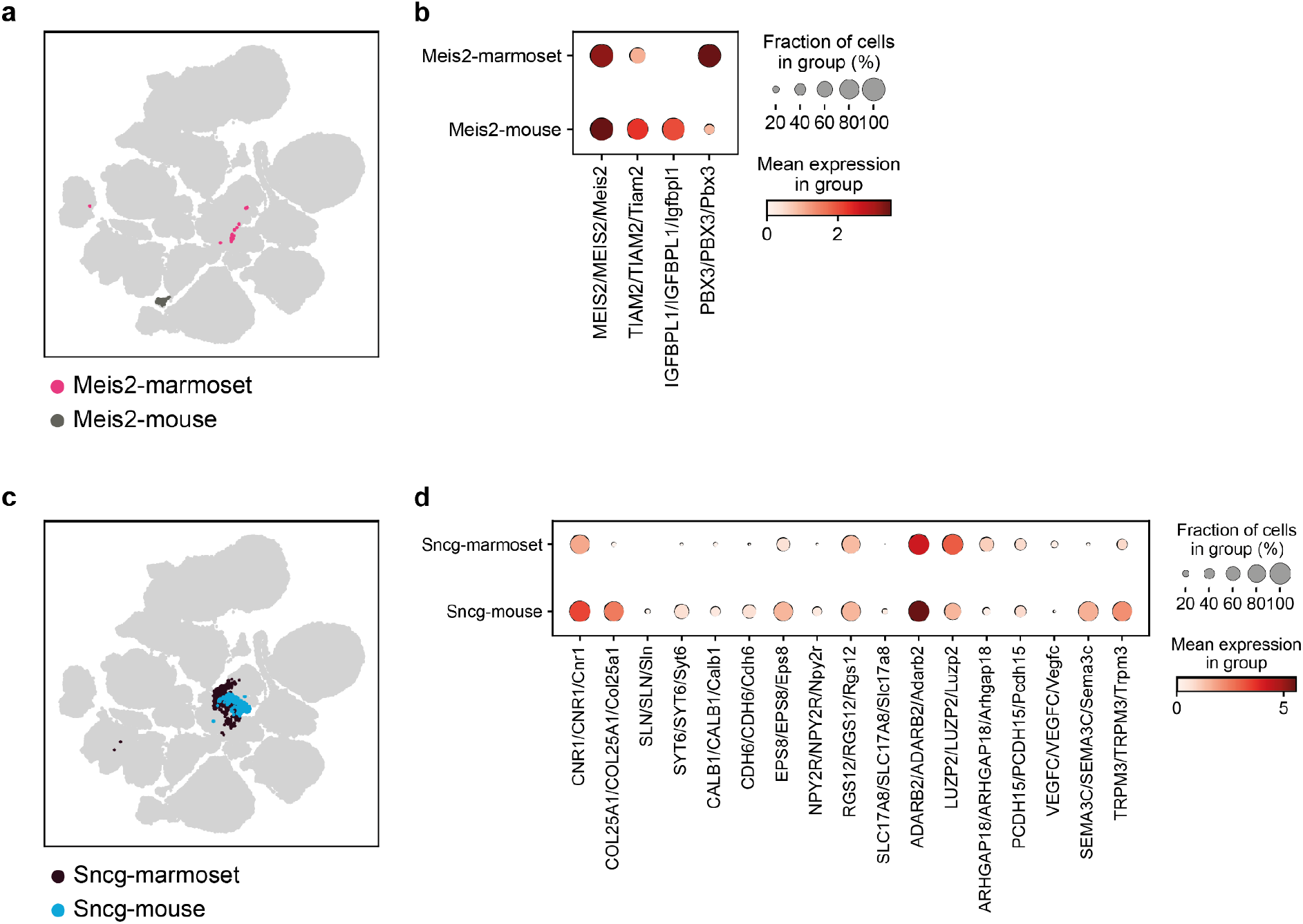
a) and c) UMAP embedding showing the integrated latent space of the reference datasets (mouse and human). The Meis2 and Sncg cell types are highlighted respectively. b) and d) Marker gene expression for the Meis2 and Sncg cell types respectively. The three gene names shown are the human/marmoset/mouse gene names.

**Figure S16:**
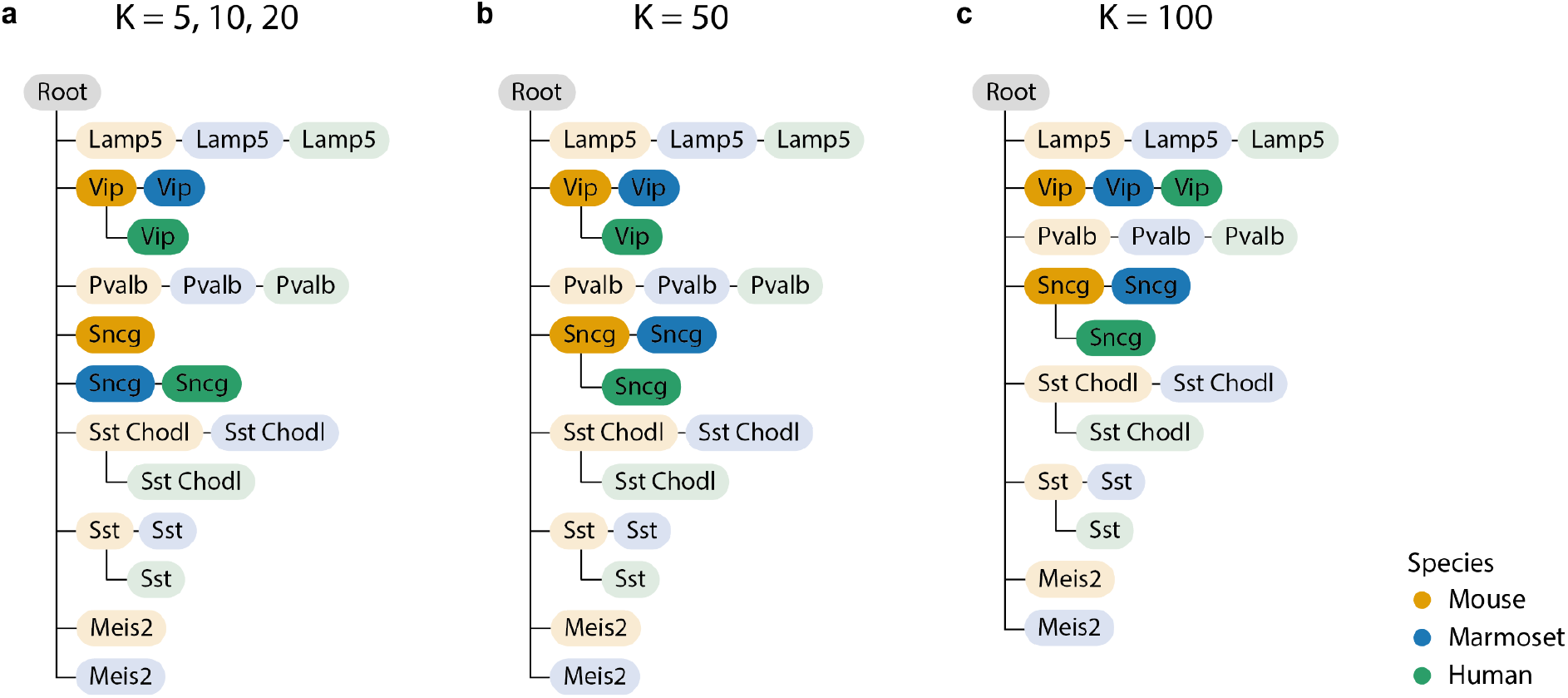
Influence of the number of neighbors (K) on the learned hierarchy. The nodes are colored according to the species they come from. The links between most nodes are robust and do not change when the number of neighbors varies. The differences between the trees are highlighted using brighter colors.

**Table S1:** Information on PBMC datasets used in this study

| Dataset (add reference) | Tissue | No. of Samples | No. of cells | No. of genes | No. of cell types | Protocol | Reference or query |
| --- | --- | --- | --- | --- | --- | --- | --- |
| Oetjen (19) | Bone marrow | 3 | 9581 | 12303 | 16 | 10X v2 | Reference |
| Sun (21) | PBMC | 4 | 8829 | 12303 | 10 | 10X | Reference |
| Freytag (20) | PBMC | 1 | 3347 | 12303 | 9 | 10X v2 | Reference |
| 10X (27) | PBMC | 1 | 10727 | 12303 | 12 | 10X v3 | Query |

**Table S2:** Overview of cell types (original labels) in the PBMC datasets

| cell type | Oetjen | Sun | Freytag | 10X |
| --- | --- | --- | --- | --- |
| CD4+ T | 2524 | 4312 | 1238 | 2937 |
| CD8+ T | 985 | 578 | 270 | 350 |
| NKT | 608 | 649 | 432 | 1056 |
| NK | 89 | 973 | 476 | 756 |
| CD20+ B | 491 | 409 | 427 | 1546 |
| CD10+ B | 207 |  |  |  |
| CD14+ MC | 997 | 1501 | 452 | 3388 |
| CD16+ MC | 165 | 271 | 25 | 364 |
| MC derived DC | 214 | 82 |  | 182 |
| pDC | 133 | 40 | 11 | 81 |
| MK progenitor | 219 | 14 | 16 | 21 |
| Erythrocytes | 1502 |  |  |  |
| Erythroid prog. | 463 |  |  |  |
| HSPC | 445 |  |  | 28 |
| MC progenitor | 428 |  |  |  |
| Plasma cell | 111 |  |  | 18 |

**Table S3:** Cell type labels of the PBMC datasets after relabeling.

| Original label | Oetjen | Sun | Freytag | 10X |
| --- | --- | --- | --- | --- |
| CD4+ T | T cells | Lymphoid | CD4+ T | CD4+ T |
| CD8+ T | T cells | Lymphoid | CD8+ T | CD8+ T |
| NKT | NKT | Lymphoid | NKT | NKT |
| NK | NK | Lymphoid | NK | NK |
| CD20+ B | CD20+ B | Lymphoid | CD20+ B | CD20+ B |
| CD10+ B | CD10+ B |  |  |  |
| CD14+ MC | MC | Myeloid | CD14+ MC | CD14+ MC |
| CD16+ MC | MC | Myeloid | CD16+ MC | CD16+ MC |
| MC derived DC | MC derived DC | Myeloid |  | MC derived DC |
| pDC | pDC | Myeloid | pDC | pDC |
| MK progenitor | MK progenitor | MK progenitor | MK progenitor | MK progenitor |
| Erythrocytes | Erythrocytes |  |  |  |
| Erythroid prog. | Erythroid prog. |  |  |  |
| HSPC | HSPC |  |  | HSPC |
| MC progenitor | MC progenitor |  |  |  |
| Plasma cell | Plasma cell |  |  | Plasma cell |

**Table S4:** Quantitative evaluation of the cell type matching algorithms.

| Method | Correct edges | Missing edges | Wrong edges |
| --- | --- | --- | --- |
| treeArches | 24 | 2 | 0 |
| FR-Match | 19 | 7 | 11 |
| MetaNeighbor | 15 | 11 | 8 |

**Table S5:** Information on brain datasets used in this study

| Species | No. of cells | No. of genes | No. of cell types<br>(class/subclass/<br>RNA_cluster) | Protocol |
| --- | --- | --- | --- | --- |
| Mouse | 159,739 | 27,439 | 3/23/116 | 10X v3 |
| Marmoset | 69,279 | 27,466 | 3/22/94 | 10X v3 |
| Human | 76,621 | 32,991 | 3/20/127 | 10X v3 |

## Notes

### Summary of Updates

Included extra experiments to emphasize that no method currently exists with the same functionality as treeArches.

## References

1. Suo, C., Dann, E., Goh, I., Jardine, L., Kleshchevnikov, V., Park, J.-E., Botting, R.A., Stephenson, E., Engelbert, J., Tuong, Z.K., et al. (2022) Mapping the developing human immune system across organs. Science, 10.1126/science.abo0510.

2. Sikkema, L., Strobl, D., Zappia, L., Madissoon, E., Markov, N.S., Zaragosi, L., Ansari, M., Arguel, M., Apperloo, L., Bécavin, C., et al. (2022) An integrated cell atlas of the human lung in health and disease. bioRxiv, 10.1101/2022.03.10.483747.

3. Tabula Sapiens Consortium*, Jones, R.C., Karkanias, J., Krasnow, M.A., Pisco, A.O., Quake, S.R., Salzman, J., Yosef, N., Bulthaup, B., Brown, P., et al. (2022) The Tabula Sapiens: A multiple-organ, single-cell transcriptomic atlas of humans. Science, 376, eabl4896.

4. Hao, Y., Hao, S., Andersen-Nissen, E., Mauck, W.M., Zheng, S., Butler, A., Lee, M.J., Wilk, A.J., Darby, C., Zager, M., et al. (2021) Integrated analysis of multimodal single-cell data. Cell, 0.

5. Swamy, V.S., Fufa, T.D., Hufnagel, R.B. and McGaughey, D.M. (2021) Building the mega single-cell transcriptome ocular meta-atlas. Gigascience, 10.

6. Osorio, D., McGrail, D.J., Sahni, N. and Stephen Yi, S. (2022) Drug combination prioritization for cancer treatment using single-cell RNA-seq based transfer learning. bioRxiv, 10.1101/2022.04.06.487357.

7. Bharat, A., Querrey, M., Markov, N.S., Kim, S., Kurihara, C., Garza-Castillon, R., Manerikar, A., Shilatifard, A., Tomic, R., Politanska, Y., et al. (2020) Lung transplantation for patients with severe COVID-19. Sci. Transl. Med., 12.

8. Wang, M., Zadeh, S., Pizzolla, A., Thia, K., Gyorki, D.E., McArthur, G.A., Scolyer, R.A., Long, G., Wilmott, J.S., Andrews, M.C., et al. (2022) Characterization of the treatment-naive immune microenvironment in melanoma with BRAF mutation. J Immunother Cancer, 10.

9. Diehl, A.D., Meehan, T.F., Bradford, Y.M., Brush, M.H., Dahdul, W.M., Dougall, D.S., He, Y., Osumi-Sutherland, D., Ruttenberg, A., Sarntivijai, S., et al. (2016) The Cell Ontology 2016: enhanced content, modularization, and ontology interoperability. J. Biomed. Semantics, 7, 44.

10. Michielsen, L., Reinders, M.J.T. and Mahfouz, A. (2021) Hierarchical progressive learning of cell identities in single-cell data. Nat. Commun., 12, 1–12.

11. Novella-Rausell, C., Grudniewska, M., Peters, D.J.M. and Mahfouz, A. (2022) A comprehensive mouse kidney atlas enables rare cell population characterization and robust marker discovery. bioRxiv, 10.1101/2022.07.02.498501.

12. Lotfollahi, M., Naghipourfar, M., Luecken, M.D., Khajavi, M., Büttner, M., Wagenstetter, M., Avsec,Ž., Gayoso, A., Yosef, N., Interlandi, M., et al. (2022) Mapping single-cell data to reference atlases by transfer learning. Nat. Biotechnol., 40, 121–130.

13. Kang, J.B., Nathan, A., Weinand, K., Zhang, F., Millard, N., Rumker, L., Moody, D.B., Korsunsky, I. and Raychaudhuri, S. (2021) Efficient and precise single-cell reference atlas mapping with Symphony. Nat. Commun., 12, 5890.

14. Gayoso, A., Lopez, R., Xing, G., Boyeau, P., Valiollah Pour Amiri, V., Hong, J., Wu, K., Jayasuriya, M., Mehlman, E., Langevin, M., et al. (2022) A Python library for probabilistic analysis of single-cell omics data. Nat. Biotechnol., 40, 163–166.

15. Lotfollahi, M., Wolf, F.A. and Theis, F.J. (8/2019) scGen predicts single-cell perturbation responses. Nat. Methods, 16, 715–721.

16. Luecken, M.D., Büttner, M., Chaichoompu, K., Danese, A., Interlandi, M., Mueller, M.F., Strobl, D.C., Zappia, L., Dugas, M., Colomé-Tatché, M., et al. (2022) Benchmarking atlas-level data integration in single-cell genomics. Nat. Methods, 19, 41–50.

17. Johnson, J., Douze, M. and Jégou, H. (2021) Billion-Scale Similarity Search with GPUs. IEEE Transactions on Big Data, 7, 535–547.

18. Pedregosa, F., Varoquaux, G., Gramfort, A., Michel, V., Thirion, B., Grisel, O., Blondel, M., Prettenhofer, P., Weiss, R., Dubourg, V., et al. (2011) Scikit-learn: Machine Learning in Python.

19. Oetjen, K.A., Lindblad, K.E., Goswami, M., Gui, G., Dagur, P.K., Lai, C., Dillon, L.W., McCoy, J.P. and Hourigan, C.S. (2018) Human bone marrow assessment by single-cell RNA sequencing, mass cytometry, and flow cytometry. JCI Insight, 3.

20. Freytag, S., Tian, L., Lönnstedt, I., Ng, M. and Bahlo, M. (2018) Comparison of clustering tools in R for medium-sized 10x Genomics single-cell RNA-sequencing data. F1000Res., 7, 1297.

21. Sun, Z., Chen, L., Xin, H., Jiang, Y., Huang, Q., Cillo, A.R., Tabib, T., Kolls, J.K., Bruno, T.C., Lafyatis, R., et al. (2019) A Bayesian mixture model for clustering droplet-based single-cell transcriptomic data from population studies. Nat. Commun., 10, 1649.

22. Bakken, T.E., Jorstad, N.L., Hu, Q., Lake, B.B., Tian, W., Kalmbach, B.E., Crow, M., Hodge, R.D., Krienen, F.M., Sorensen, S.A., et al. (2021) Comparative cellular analysis of motor cortex in human, marmoset and mouse. Nature, 598, 111–119.

23. Zhang, Y., Aevermann, B., Gala, R. and Scheuermann, R.H. (2022) Cell type matching in single-cell RNA-sequencing data using FR-Match. Sci. Rep., 12, 9996.

24. Zhang, Y., Aevermann, B.D., Bakken, T.E., Miller, J.A., Hodge, R.D., Lein, E.S. and Scheuermann, R.H. (2021) FR-Match: robust matching of cell type clusters from single cell RNA sequencing data using the Friedman-Rafsky non-parametric test. Brief. Bioinform., 22.

25. Aevermann, B., Zhang, Y., Novotny, M., Keshk, M., Bakken, T., Miller, J., Hodge, R., Lelieveldt, B., Lein, E. and Scheuermann, R.H. (2021) A machine learning method for the discovery of minimum marker gene combinations for cell type identification from single-cell RNA sequencing. Genome Res., 31, 1767–1780.

26. Crow, M., Paul, A., Ballouz, S., Huang, Z.J. and Gillis, J. (2018) Characterizing the replicability of cell types defined by single cell RNA-sequencing data using MetaNeighbor. Nat. Commun., 9, 884.

27. Genomics,10x (2018) 10x Datasets Single Cell Gene Expression.

28. Abdelaal, T., Michielsen, L., Cats, D., Hoogduin, D., Mei, H., Reinders, M.J.T. and Mahfouz, A. (2019) A comparison of automatic cell identification methods for single-cell RNA sequencing data. Genome Biol., 20, 194.

29. Madissoon, E., Oliver, A.J., Kleshchevnikov, V., Wilbrey-Clark, A., Polanski, K., Orsi, A.R., Mamanova, L., Bolt, L., Richoz, N., Elmentaite, R., et al. (2021) A spatial multi-omics atlas of the human lung reveals a novel immune cell survival niche. bioRxiv, 10.1101/2021.11.26.470108.

30. Basil, M.C., Cardenas-Diaz, F.L., Kathiriya, J.J., Morley, M.P., Carl, J., Brumwell, A.N., Katzen, J., Slovik, K.J., Babu, A., Zhou, S., et al. (2022) Human distal airways contain a multipotent secretory cell that can regenerate alveoli. Nature, 604, 120–126.

31. Kadur Lakshminarasimha Murthy, P., Sontake, V., Tata, A., Kobayashi, Y., Macadlo, L., Okuda, K., Conchola, A.S., Nakano, S., Gregory, S., Miller, L.A., et al. (2022) Human distal lung maps and lineage hierarchies reveal a bipotent progenitor. Nature, 604, 111–119.

32. Rustam, S., Hu, Y., Mahjour, S.B., Rendeiro, A.F., Ravichandran, H., Urso, A., D’Ovidio, F., Martinez, F.J., Altorki, N.K., Richmond, B., et al. (2023) A Unique Cellular Organization of Human Distal Airways and Its Disarray in Chronic Obstructive Pulmonary Disease. Am. J. Respir. Crit. Care Med., 10.1164/rccm.202207-1384OC.

33. Tsukui, T., Sun, K.-H., Wetter, J.B., Wilson-Kanamori, J.R., Hazelwood, L.A., Henderson, N.C., Adams, T.S., Schupp, J.C., Poli, S.D., Rosas, I.O., et al. (2020) Collagen-producing lung cell atlas identifies multiple subsets with distinct localization and relevance to fibrosis. Nat. Commun., 11, 1920.

34. Morse, C., Tabib, T., Sembrat, J., Buschur, K.L., Bittar, H.T., Valenzi, E., Jiang, Y., Kass, D.J., Gibson, K., Chen, W., et al. (2019) Proliferating SPP1/MERTK-expressing macrophages in idiopathic pulmonary fibrosis. Eur. Respir. J., 54.

35. Karman, J., Wang, J., Bodea, C., Cao, S. and Levesque, M.C. (2021) Lung gene expression and single cell analyses reveal two subsets of idiopathic pulmonary fibrosis (IPF) patients associated with different pathogenic mechanisms. PLoS One, 16, e0248889.

